# XSSDense: X-ray Solution Scattering Density Reconstruction Using a Variational Autoencoder

**DOI:** 10.64898/2026.08.07.743437

**Authors:** Leonardo Monrroy, Sebastian Cardoch, Seoyoung Lee, Sebastian Westenhoff

## Abstract

Solution X-ray scattering reveals the structure of biomolecules in solution. Scattering profiles also encode information on conformational heterogeneity and, when measured with time resolution, transient structural intermediates. However, this information is difficult to access because scattering profiles contain limited information, and their interpretation largely relies on fitting candidate structures guided by prior knowledge. Direct reconstruction of electron densities has remained challenging because of the difficulty of incorporating such priors. Here we introduce XSSDense, which combines a variational autoencoder trained on electron densities from predicted or modelled protein ensembles with genetic optimization to refine densities against scattering data. We verify the method on synthetic data and apply it to time-resolved scattering experiments, recovering the conformational heterogeneity of the unfolded state of *As*LOV2, resolving a de novo density of its pre-unfolding intermediate, and characterizing the signalling-state ensemble of photoactive yellow protein. We further demonstrate its application to conventional SAXS by refining different conformations of adenylate kinase. XSSDense enables structurally grounded electron density reconstructions that intrinsically capture conformational heterogeneity.

## Main

Proteins and nucleic acids rely on structural changes for their biological function. Lig- and binding, enzyme catalysis, signal transduction, molecular recognition, and nucleic acid folding all involve transitions between distinct conformational states. Under bio-logically relevant conditions, these macromolecules are intrinsically dynamic and often populate ensembles of interconverting conformations rather than single well-defined structures [1]. Characterizing both the conformational transitions and the underlying conformational heterogeneity is therefore essential for understanding biological function.

Small- and wide-angle X-ray scattering (S/WAXS) is particularly well suited to study these processes because it probes biomolecular structure directly in solution at room temperature. For example, it is used to investigate solution structures of biomolecules [2], discover novel biophysics [3], and provide unique input for integrative structural biology [4]. Time-resolved S/WAXS can access a broad range of timescales, enabling the study of conformational transitions in proteins [5–11], and structural dynamics of nucleic acids [12].

Because the measured S/WAXS signal represents an average over all molecular orientations and populated conformations, the resulting one-dimensional scattering profile contains only limited structural information. Nevertheless, several global structural parameters, including the radius of gyration, molecular weight, maximum particle dimension, and pair-distance distribution function, can be determined directly in a model-free manner [2]. Recovering higher-resolution information, however, is considerably more challenging.

Obtaining more precise structural information requires the incorporation of external structural priors, for example from complementary structural methods, molecular simulations, or protein structure prediction [13–18]. Because these priors are naturally represented in atomic-coordinate space, they have almost exclusively been incorporated into S/WAXS analysis through *forward* modelling (Fig. 1a, middle), in which chemically plausible candidate structures are simulated, predicted, or otherwise determined and subsequently evaluated against the experimental scattering data [6, 9, 11, 15, 19–21]. While highly successful, forward modelling ultimately refines one or a small number of discrete structural models. Consequently, it is prone to overfitting and struggles to represent the broad conformational heterogeneity commonly expected under physiological conditions. Although increasingly sophisticated ensemble refinement methods based on molecular dynamics and Bayesian or maximum entropy approaches have been developed [17, 18, 22], faithfully representing broad conformational distributions remains challenging.

**Figure 1.**
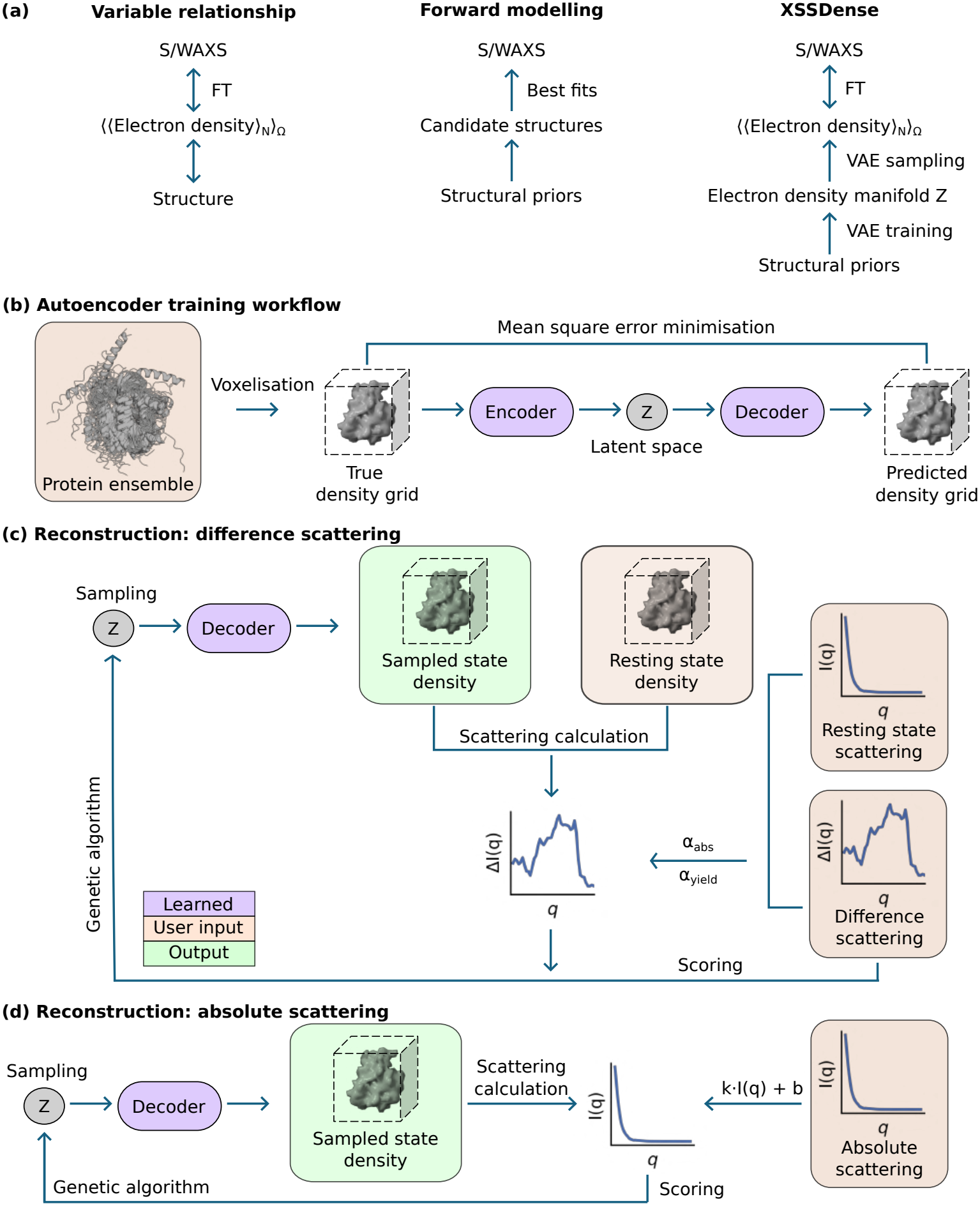
(a) Conceptual idea behind XSSDense. (left) An experimentally measured scattering intensity corresponds to an orientation-(Ω) and ensemble-(*N* ) averaged electron density in solution. (middle) In forward-modelling approaches, candidate structures representing a single or limited set of states are compared to S/WAXS data. (right) XSSDense learns chemical and structural constraints from supplied structural ensembles and then generates electron densities that fit the S/WAXS patterns. (b) Training data from AFsample2 [37] or molecular dynamics simulations are converted into electron density maps using atomic form factors. A variational autoencoder (VAE) encodes densities into a compressed representation *z* and decodes them back by minimizing the error between the input and output. (c) The trained latent space is explored for possible conformations which, after subtraction of the resting-state, give a difference scattering signal. The difference scattering is scaled to experimental units, compared with the experimental data, and ranked accordingly. A new round of sampling is then guided by the genetic algorithm until convergence. (d) Absolute scattering can also be fitted if desired. Here, *k* and *b* in *kI*(*q*) + *b* are determined by *q*-weighted least squares in each iteration.

The measured scattering intensity is related to the squared modulus of the Fourier transform of the electron-density distribution of the sample (Fig. 1a, left). Because electron densities naturally represent conformational heterogeneity through regions of reduced density arising from fractional occupancies [23], they provide a more natural representation than discrete atomic-coordinate models. A natural solution would therefore be to refine three-dimensional electron densities directly from the scattering data. However, recovering electron densities directly from the measured scattering profile constitutes an ill-posed inverse problem. Direct reconstructions of electron density or dummy-atom models remain fundamentally limited to resolutions of only tens of ångströms [24]. Even iterative phase-recovery approaches for non-uniform density reconstruction provide only modest improvements in resolving power [25]. Achieving higher resolution and greater structural specificity therefore requires the incorporation of structural priors into the electron-density refinement. Since atomic coordinates are no longer represented explicitly, however, incorporating structural and physical priors into electron-density refinement has so far remained prohibitively challenging.

Here, we address this challenge by learning a physically accessible electron-density manifold from an ensemble of candidate structures. The manifold acts as a structural prior during refinement, allowing three-dimensional electron density maps to be reconstructed directly from S/WAXS data, while remaining consistent with physically plausible protein conformations (Fig. 1a, right). Conformational flexibility is naturally encoded in the spatial distribution and magnitude of the electron density.

We hypothesize that a variational autoencoder (VAE) can learn a manifold of physically accessible electron densities from an ensemble of candidate structures [26]. VAEs are known to learn continuous, low-dimensional representations of complex structural ensembles [27–32]. We further hypothesize that the resulting continuous probabilistic latent space enables efficient interpolation between physically plausible electron densities, thereby capturing conformational heterogeneity. Refinement within this electron-density manifold, constrained by the experimental S/WAXS data, should therefore allow direct reconstruction of three-dimensional electron densities while remaining consistent with physically realistic protein conformations. Our work follows a general trend in which generative models are becoming increasingly prominent in molecular design and structure prediction [33–36].

We implement this framework in XSSDense, first validating it on synthetic crambin ensembles and then applying it to experimental time-resolved S/WAXS data from *Avena sativa* Light-Oxygen-Voltage Sensing Domain 2 (*As*LOV2) [9] and photoactive yellow protein (PYP) [8]. XSSDense reproduces the scattering data while resolving conformational heterogeneity beyond previous coordinate-based analyses. Finally, using synthetic SAXS data for adenylate kinase (AdK), we demonstrate the recovery of distinct conformational states and their populations in mixtures.

### XSSDense

Structure refinement in XSSDense consists of two steps. In the first step, a variational autoencoder (VAE) is trained on three-dimensional electron densities on a grid generated from ensembles of simulated or predicted protein structures (Fig. 1b). The VAE compresses each density into a low-dimensional latent representation (encoder) and reconstructs it back into a three-dimensional density (decoder). Training minimizes the difference between the original and reconstructed densities (see Methods). In the second step, the method can either be used to fit difference or absolute scattering, such as in time-resolved X-ray scattering or conventional SAXS experiments, respectively (Fig. 1c and d). In both cases, the trained latent space is sampled to generate electron densities that fit an experimental X-ray scattering profile. The refinement is done by a genetic algorithm [38], and candidate solutions are ranked according to the *R*^2^ (Equ. (13)) between the observed and calculated scattering profiles. The latent space is iteratively resampled around the highest-scoring solutions to generate improved candidate densities. Reconstructed densities are low-pass filtered at 0.6*/*Å, comparable to the highest *q* value used in scoring, to suppress unsupported high-resolution features (see Methods). Calculated scattering profiles include contributions from displaced solvent and the hydration layer, which are important to consider for solution scattering (see Methods). For absolute scattering, the scale between experiment and theory is treated as a nuisance parameter and determined by least squares at each refinement iteration (Fig. 1d), but for difference scattering the reconstructions are restrained to match a target photoactivation yield, which is relevant for correctly estimating the degree of structural change (Fig. 1c) [9].

### Validation on synthetic data

First, we tested XSSDense using a synthetic test case. Crambin is a small protein (4.74 kDa) with three sulphur bridges and a secondary structure consisting of two *α* helices, a *β* sheet, and flexible loops. A molecular dynamics simulation of a single crambin molecule (PDB ID 1CRN) [39] revealed that the *α* helices and connecting loop remained rigid, whereas the *β* sheet and terminal loop partially unfolded (Fig. 2b inset).

**Figure 2.**
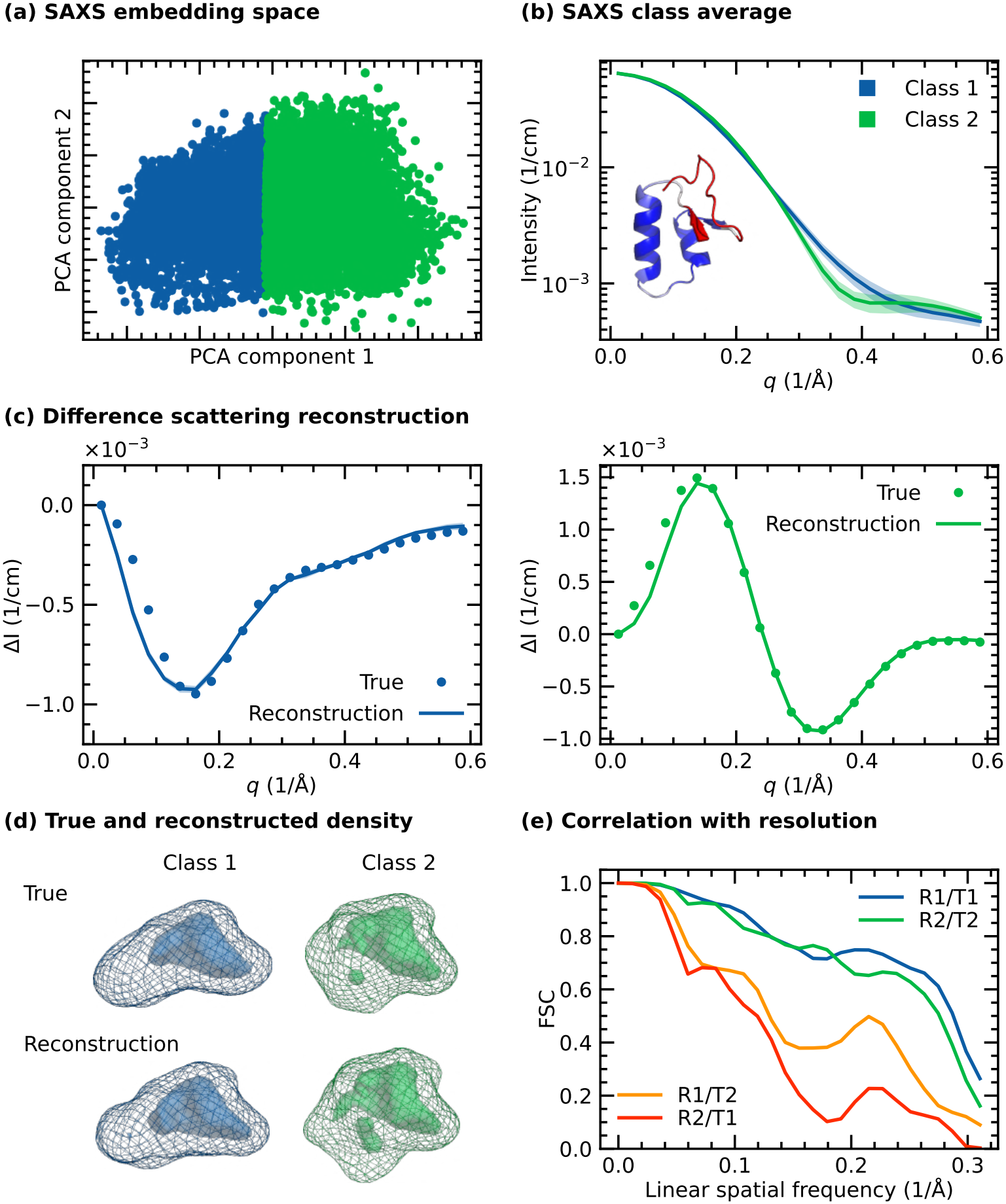
(a) Lower-dimensional representation and clustering of *in vacuo* small-angle X-ray scattering (SAXS) profiles of crambin obtained from molecular dynamics simulations. (b) Ensemble-average SAXS profiles (solid line) and standard deviation (shaded region) for the two identified classes. Inset shows crambin’s reference structure as a cartoon representation colour-coded to show regions with low (blue) and high (red) root mean square fluctuation based on molecular dynamics simulations. The *α* helices and loop connecting them are rigid, while the *β* sheet and terminal loop are flexible. (c) Comparing true and reconstructed ensemble-average difference scattering curves over 10 trials for the two classes as a function of momentum transfer *q* = 4*π* sin(*θ*)*/λ*. (d) Comparing true and reconstructed ensemble average electron densities plotted as a mesh at 0.1 e*/*Å^3^ and surface at 0.3 e*/*Å^3^. The maps were subjected to a low-pass Gaussian filter with a half-power point of 0.6*/*Å. (e) Fourier shell correlation (FSC) between low-pass filtered true (T) and reconstructed (R) electron densities between the different classes.

For each trajectory frame we computed S/WAXS profiles of the protein in vacuum up to *q* = 0.6*/*Å (see Methods). Principal component analysis and k-means clustering (Fig. S1) revealed that two components explained more than 95 % of the variance in the SAXS curves (Fig. 2a). We separated the embedding space into two groups, each representing an ensemble of related protein conformations sampled during the simulation, resulting in class-averaged SAXS profiles that differed beyond the variability within each group (Fig. 2b). The corresponding difference scattering was generated by subtracting the scattering calculated from the VAE-reconstructed electron density of the first molecular-dynamics frame from the class-averaged SAXS profiles. This was done to remove the systematic error introduced by the VAE in the density reconstruction. Averages over the electron densities of the two ensembles indicate two distinct conformational states, with class 1 exhibiting a more elongated conformation and class 2 a more globular shape (Fig. 2d).

Following the XSSDense workflow, we first trained the variational autoencoder on the electron densities of each simulation frame. Evaluation using electron densities from an independent test dataset not used during training indicates high reconstruction accuracy by the autoencoder (Fig. S2a). In the second step, we reconstructed the class-averaged electron densities by specifying the corresponding class-averaged SAXS profiles as refinement targets. The reconstruction algorithm converged after approximately 10 iterations, with score values above 0.95 (Fig. S2b). The reconstructed curve shows good agreement with the true difference scattering (Fig. 2c), and the reconstructed densities correctly recover the defining features for both classes (Fig. 2d).

As in typical experimental data, the finite *q*-range in this synthetic example limits the spatial resolution that can be constrained during refinement. Consequently, we initially observed high-resolution features in the reconstructed electron densities that are absent from the true densities (Fig. S3). To suppress them, we applied the lowpass filter described above (half-power point at 0.6*/*Å; see Methods), yielding densities whose information content is consistent with both the S/WAXS data and the learned electron-density manifold (Fig. S3).

To quantify the reconstruction accuracy, we computed the Fourier shell correlation (FSC) between the true and reconstructed electron densities (Fig. 2e). The FSC remains high (*>*0.5) up to a spatial frequency of approximately 0.3*/*Å (3.3 Å resolution; 1*/d*), demonstrating high structural fidelity. Consistent with this result, the Pearson correlation coefficient between the reconstructed and true densities approaches 1 at lower resolution (Fig. S4). Interestingly, this resolution exceeds the nominal resolution range of the data, which extends to *q* = 0.6*/*Å (10.5 Å; *q* = 2*π/d*). In contrast, the FSC with the true density of the opposing class drops rapidly, confirming that XSSDense correctly captures the structural features that distinguish the two conformational ensembles.

Overall, this synthetic example demonstrates that XSSDense can accurately recover ensemble-averaged electron densities directly from difference SAXS profiles, producing reconstructions with remarkably high structural fidelity.

### Solving the unfolded state of *As*LOV2

Having established the functionality of the method, we now apply it to an experimental dataset. *As*LOV2 is a blue-light receptor that uses a flavin mononucleotide (FMN) cofactor for light sensing (Fig. 3a). Upon blue-light illumination, FMN forms a covalent bond with a nearby conserved cysteine residue within microseconds [40, 41]. A larger conformational change in the protein follows, during which the structured C-terminal J*α* helix unfolds and becomes disordered [9, 42–46]. The unfolding was proposed based on nuclear magnetic resonance (NMR) spectroscopy more than two decades ago [42] and later confirmed by infrared [43, 46] and transient grating spectroscopy [44]. However, the degree of unfolding and the conformational ensemble could not be determined. Owing to their conformational response to light, LOV domains have been widely adopted in optogenetics and protein engineering [47].

**Figure 3.**
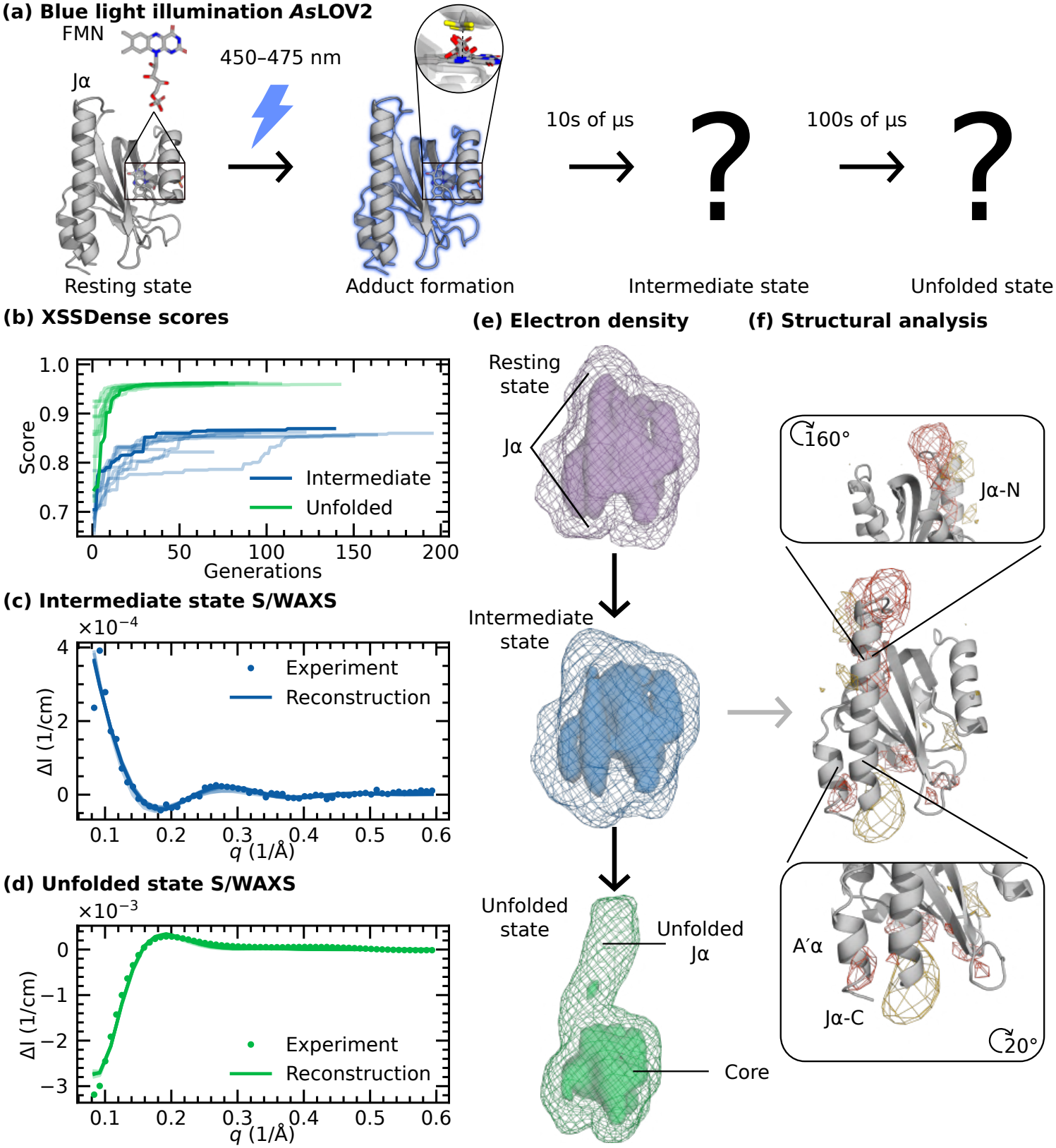
(a) Simplified photocycle of *As*LOV2 presented by Konold et al. (2024) [9]. The left panel shows the dark-adapted *As*LOV2 resting structure (PDB ID 7PGX) together with its flavin mononucleotide (FMN) cofactor. Upon blue light illumination a cysteine adduct forms within ∼2 µs leading to the emergence of an intermediate state followed by the unfolding of the J*α* helix. (b) XSSDense fitting score as a function of the number of iterations for 10 independent runs using random seeds, demonstrating convergence of the results. Darkest line represents the run with the highest final fitting score (b–d). (c,d) Experimental (dots) and reconstructed (line) difference scattering for the intermediate and unfolded state, respectively. (e) Voxelised density of the resting, intermediate, and unfolded states (the latter two reconstructed from the highest scoring runs shown in panels c and d). The outer mesh and inner surface are drawn at 0.05 e*/*Å^3^ and 0.3 e*/*Å^3^, respectively. (f) Low-pass filtered electron density difference map (intermediate state minus resting state) overlaid on the resting structure showing positive (gold) and negative (red) changes evaluated at *±*0.08 e*/*Å^3^ corresponding to 2*σ* level. Magnified views reveal distinct structural changes localized around the N-terminal region of the J*α* helix and around the C-terminal of J*α* helix and adjacent A’*α* helix. The intermediate density is scaled to match the total resting density prior to subtraction.

Two recent time-resolved S/WAXS studies have attempted to resolve the conformational ensemble of the disordered state [9, 11]. Both detected a microsecond rise in the difference S/WAXS signal and interpreted it as the unfolded state using forward modelling. Since this relied on refining *single structures* to a signal that essentially reflects an average over a broad *structural ensemble*, the structural interpretation may be biased. Determining ensemble-averaged electron densities, as enabled by XSS-Dense, provides a more appropriate framework. Here we used difference scattering corresponding to the kinetically resolved unfolded state [9].

To reconstruct the electron density of the unfolded state, we generated 10 000 *As*LOV2 structures using AFsample2, which employs multiple sequence alignment masking to expand the conformational space sampled by AlphaFold2 [37]. These structures were used to train the VAE implemented in XSSDense (Methods and Fig. S5a). We used a strongly regularised latent space (*β* = 1), restricting the search to conformations close to the training ensemble and thereby reducing the risk of overfitting the limited Δ*I* information. During reconstruction, scattering contributions from vacuum, bulk solvent, and hydration shell were included (Methods and Fig. S6). The hydration-shell thickness and density contrast were determined from the resting state within physically motivated bounds (see Methods). The optimisation yielded values of 5.0 Å and 0.043 e*/*Å^3^, respectively (Tab. S1), corresponding to the upper bounds of the allowed parameter range. The excitation yield was constrained to the experimentally determined value of approximately 15 % [9]. To assess the robustness of the reconstruction, we performed 10 independent runs with different seeds. All runs converged to a final score larger than 0.8 (Fig. 3b).

The reconstructions show an excellent fit to the time-resolved S/WAXS difference scattering signal of the intermediate and unfolded states (Fig. 3c,d). Across the ten runs, the optimized excitation yields were 11(3) % and 16.0(3) % for the intermediate and unfolded states, respectively. We manually varied the hydration-shell parameters and found that the unfolded state yield was largely unaffected by the hydration-shell choice, while the intermediate state yield varied between 5 % to 19 % and should therefore be considered approximate.

The resulting density of the unfolded state (green) shows clear characteristics of unfolding when compared to the density of the resting state (magenta) (Fig. 3e). The extent of the core (*>*0.3 e*/*Å^3^) decreases relative to the resting state and a new extended part with a lower density (∼0.1 e*/*Å^3^) emerges. The reduced electron density reflects the expected conformational heterogeneity of the unfolded helix and indicates that XSSDense interpolates smoothly across structures. This interpretation is supported by the observation that for the 10 training densities closest to the reconstructed density, the FSC remains equally high (Fig. S8), which demonstrates that XSSDense identifies a region of conformational space rather than selecting a single electron density from the learned manifold. Overall, these results provide confidence that conformationally heterogeneous states can now be rigorously characterized by X-ray solution scattering.

### Revealing the structure of a new, pre-unfolding state in *As*LOV2

In addition to the unfolded state, a time-resolved S/WAXS study also identified an intermediate state populated between adduct formation and J*α* unfolding [9]. However, because its time-resolved S/WAXS signal was found to be very small, its structure could not be resolved by single-structure *forward* modelling alone [9]. Molecular dynamics (MD) simulations and time-resolved infrared (IR) spectroscopy have predicted the existence of one or more such intermediate states [46]. Resolving the structure of this state is important to gain mechanistic insight into how the helix unfolding is triggered in LOV2 domains.

Here, we used the S/WAXS profile of the intermediate state as input to XSS-Dense, which reconstructed an electron density whose calculated scattering profile closely reproduced the experimental data (Fig. 3c). Consistent with the weak scattering signal, the reconstructed averaged electron density (blue) differs only subtly from the reference state (Fig. 3e). To visualize these changes more clearly, we subtracted the electron density of the reference state that had been encoded/decoded beforehand (Fig. 3f). The resulting difference density is localized primarily to the N- and C-terminal ends of the J*α* helix, with the C-terminal changes extending into the A’*α* helix. The central part of the J*α* helix remains largely unchanged. This contrasts with molecular dynamics simulations, which predicted a partially detached J*α* helix in the intermediate state [46]. Although the reconstructed electron density does not yet provide atomic-resolution structural information, it identifies the A’*α* helix as a key structural element in preparing J*α*-helix unfolding, thereby substantially narrowing the search for the molecular origin of this transition.

### Capturing N-terminal rearrangements in PYP

As a third experimental case, we investigate photoactive yellow protein (PYP), involved in phototaxis from *Halorhodospira halophila* bacteria [48]. It contains a paracoumaric acid (pCA) bound via a conserved cysteine as a light-sensitive chromophore (Fig. 4a). Its photocycle has been actively investigated using simulations and time-resolved experimental methods [8, 15, 49–52]. After photoexcitation by blue light, the chromophore undergoes trans-cis isomerisation within picoseconds [50], and the reaction proceeds through several intermediate states before reaching the final signalling state (pB1) [49, 53]. While it is clear that the N terminus unfolds [54], the extent of this process and its structural and dynamical consequences remain highly debated.

**Figure 4.**
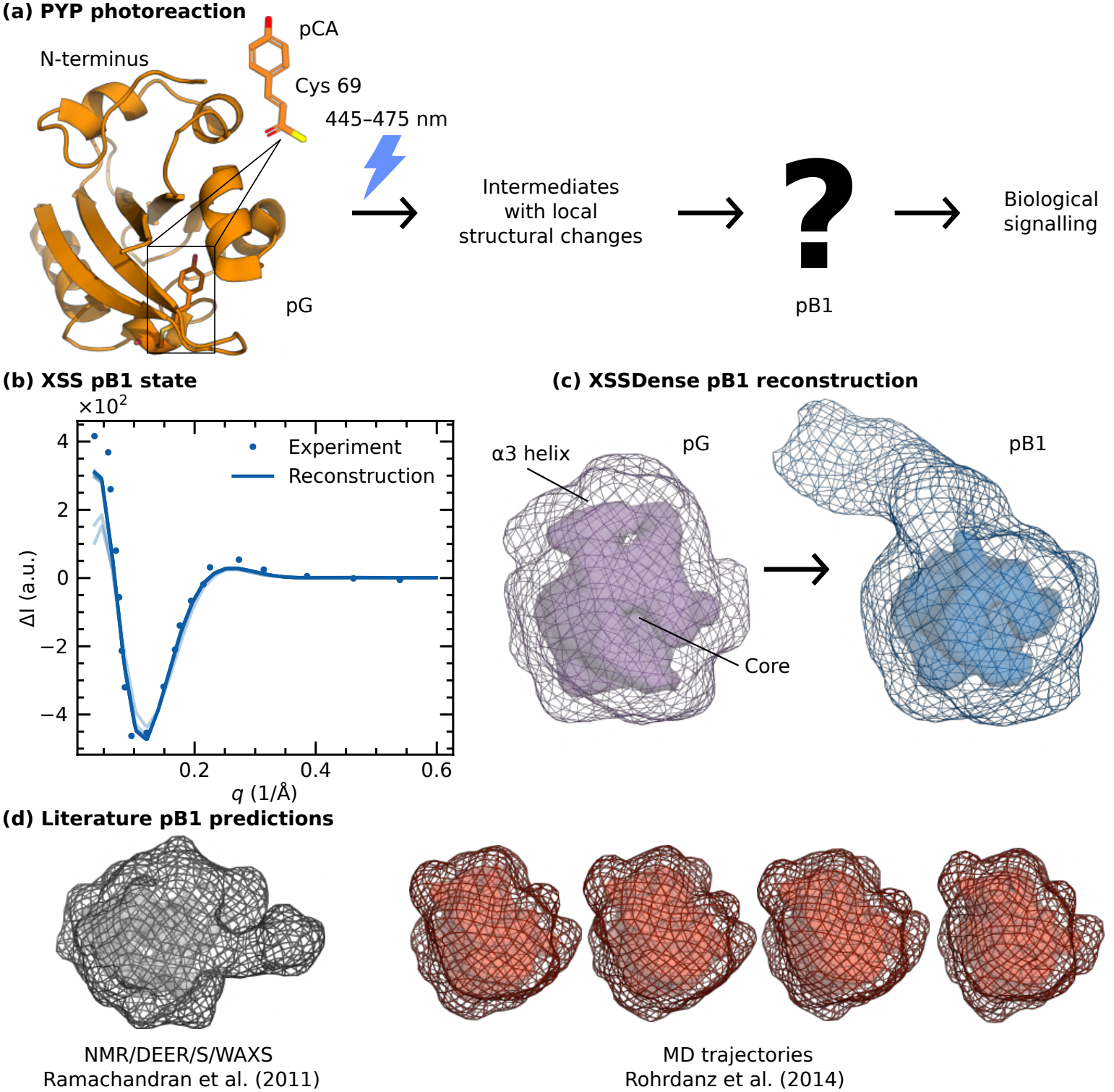
(a) Simplified diagram of PYP’s photoreaction cycle. The dark-adapted photoactive yellow protein (PDB ID 6P5G) has its pCA chromophore bound via a conserved cysteine residue [56]. Following photoexcitation and several intermediate states, the signalling state, pB1, is reached within milliseconds. Here, a large conformational change takes place, leading to biological signalling. (b) Comparison of the experimental difference scattering curve (dots) for the pB1 signalling state extracted from Cho et al. (2016) [8] and the reconstructed scattering curve as produced by XSSDense averaged over 10 trials (solid line) along with the standard deviation (shaded region). (c) Electron density of the resting-state pG (purple) and the reconstructed density of the pB1 state averaged over the 10 trials (blue). (d) Density reconstructions from NMR/DEER/S/WAXS [15] and MD trajectories [52]. Electron density outer meshes and inner surfaces are drawn at 0.05 e*/*Å^3^ and 0.3 e*/*Å^3^, respectively.

Structural ensembles have been predicted by MD simulations [52, 55] or obtained by refining the populations of NMR- and double electron-electron resonance (DEER)-derived conformers against signalling-state S/WAXS data [13, 15]. However, these ensembles are in disagreement with each other [8, 15, 52], which may in part be due to the difficulties of refining structural heterogeneity from discrete conformational ensembles. S/WAXS measurements have revealed an increased radius of gyration and an elongated conformation [8, 15, 54]. Low-resolution envelopes have been reconstructed from the scattering data [8], but do not reveal structural details beyond the fact that the protein is somewhat elongated. Thus, it remains uncertain how many residues at the N terminus participate in the detachment process and how much structure is retained. Here, we used XSSDense to obtain additional structural information at the level of electron densities.

We first trained the XSSDense variational autoencoder on 12 500 structures from AFsample2 (Fig. S5b) and then performed 10 independent fitting trials, obtaining a converged solution in every case. The resulting difference scattering profiles closely reproduce the kinetically resolved experimental data of pB1 (Fig. 4b, extracted from [8]), demonstrating the robustness and reproducibility of our method. Across all 10 independent fitting trials, we obtained similar solutions with a fitting score of 0.88 and an estimated photoactivation yield of 46 %, which is close to the experimentally determined value of 42 % [8]. The reconstructed electron density, averaged across all trials, exhibits a compact core and a weaker and more extended density near the N terminus (Fig. 4c) that indicates a partial disordering of the protein relative to the resting state. In comparison to previous low-resolution shape reconstructions of the pB1 signalling state [8], the XSSDense reconstruction localizes the structural change to the N-terminal region and provides finer-resolution information.

We compared the reconstructed electron densities against structural ensembles derived from combined NMR/DEER/S/WAXS refinement [15] and molecular dynamics simulations [52] (Fig. 4d). The experimentally refined ensemble suggested substantial unfolding and remodelling of the core of the protein, whereas the MD-derived ensemble predicted only minor structural changes of core and *α*3 tail. Our reconstruction agrees with the NMR/DEER/S/WAXS-derived ensemble in the magnitude of the conformational change, but not in the direction of the detached *α*3 helix and several other secondary structural elements. This discrepancy may reflect the limited number of long-range distance restraints in the NMR data, potentially biasing the structural ensemble used in the previous S/WAXS refinement. In contrast, the MD-derived ensemble confines the changes to the N-terminal region, consistent with our reconstruction, but substantially underestimates their magnitude. In summary, the electron densities obtained by XSSDense suggest that the N-terminal region is indeed the focal point of the conformational change, but that the N-terminal *α*3 helix is partially unfolded.

### Recovering open and closed conformations of adenylate kinase

To test whether XSSDense can refine densities against SAXS data, and whether it can distinguish two distinct conformational states from these data, we turned to adenylate kinase (AdK) from *Escherichia coli*. AdK is a phosphotransferase (23.6 kDa) consisting of a rigid CORE domain and two mobile domains, LID and NMP, which close over the CORE upon substrate binding (Fig. 5a) [57, 58]. Its open (PDB ID 4AKE) [57] and closed (PDB ID 1AKE) [59] crystal structures provide a well-defined pair of distinct conformations, and the protein has served as a precedent for elucidating the closing mechanism using time-resolved X-ray scattering [6].

**Figure 5.**
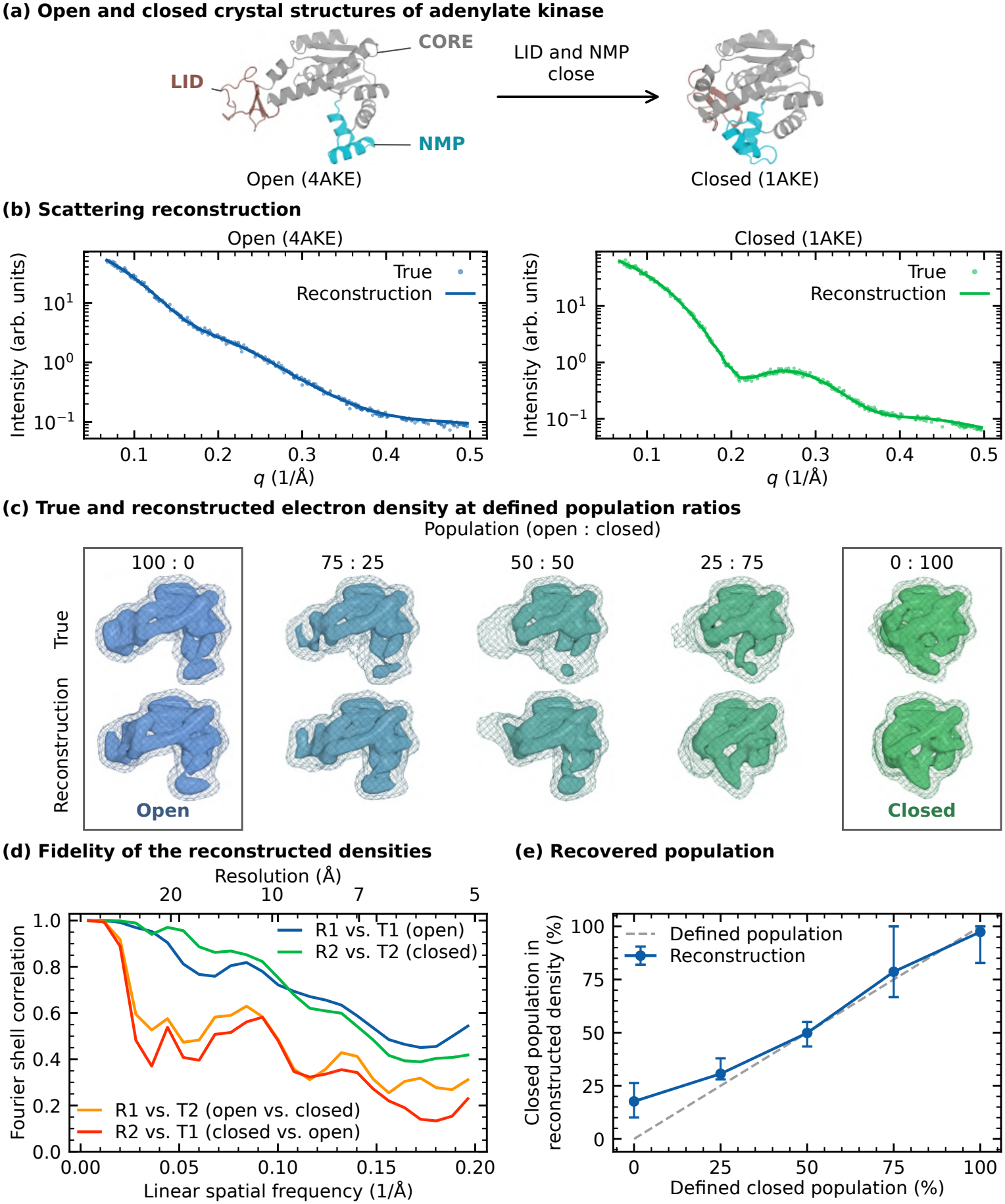
(a) Open (PDB ID 4AKE) and closed (PDB ID 1AKE) crystal structures of adenylate kinase (AdK) with the LID (brown), NMP (teal) and CORE (grey) domains. (b) Comparing true (dots) and reconstructed (line) scattering curves for the open and closed states as a function of momentum transfer *q* = 4*π* sin(*θ*)*/λ*. Reconstructed curves are averaged over 10 trials. (c) True and reconstructed electron densities at defined population ratios of the open and closed states (open : closed), averaged over 10 trials, plotted as a mesh at 0.1 e*/*Å^3^ and surface at 0.3 e*/*Å^3^. The true densities are the population-weighted average of the densities of the crystal structures. The maps were subjected to a low-pass Gaussian filter with a half-power point of 0.6*/*Å. The pure open and closed states are boxed. (d) Fidelity of the reconstructed densities, given by the Fourier shell correlation (FSC) between low-pass filtered true (T) and reconstructed (R) electron densities of the open (1) and closed (2) states. The top axis gives the corresponding real-space resolution. (e) Closed population recovered from the reconstructed density versus the defined closed population. Points show the median and bars the range of 10 trials. The dashed line marks ideal recovery.

The variational autoencoder was trained on electron densities of 5370 structures from published molecular dynamics simulations covering both states [60, 61]. Principal component analysis and k-means clustering of their *in vacuo* SAXS profiles separated the ensemble into an open-like and a closed-like class, with the two crystal structures, which were not part of the training set, at opposite ends of the embedding space (Fig. S10).

We used SAXS profiles calculated from the two crystal structures as refinement targets, performing 10 independent fits for each using the absolute-scattering reconstruction scheme (Methods and Fig. 1d). The reconstructed curves closely reproduce both targets (*R*^2^ = 0.995; Fig. 5b). All 10 trials converged to the correct state (Fig. S11), and the averaged densities recover the characteristic domain arrangement (Fig. 5c, outer columns): the LID and NMP domains extend away from the CORE in the open reconstruction and pack onto it in the closed one.

The FSC confirms that both states are recovered with resolutions of approximately 6.5 Å (open) and 6.8 Å (closed), with substantially lower correlations to the opposing state (Fig. 5d). As for crambin, these FSC profiles indicate that the resolution of the reconstruction exceeds that of the target scattering (0.5*/*Å, 12.6 Å), indicating that the structural prior supplies features that the scattering profile alone does not constrain. Finally, we tested how a single density reconstructed by XSSDense reflects potential mixtures of the open and closed states. Target SAXS profiles were calculated as population-weighted sums of the open and closed profiles at defined ratios. These were used to reconstruct electron densities using XSSDense, which gradually changed from an open to a closed shape, with good visual agreement with the expected (true) densities (Fig. 5c). We then decomposed each reconstructed density into contributions from the open and closed crystal densities (see Methods). The obtained populations indicate that the fits recover the trend (median of 10 trials; Fig. 5e).

We find excellent agreement, but note that reconstructing a mixture of states as a single average electron density neglects the covariance between the structures of the two states (see Discussion). Since we know the ground truth in the present case, we could compute the error introduced by this approximation in the context of the fitting procedure and found it to be below 4 % (Fig. S12). This error is relatively small, particularly considering the large structural change involved. Nevertheless, the populations recovered from a single reconstructed density should not be over-interpreted when multiple states are present.

Overall, given a structural prior encompassing both states, XSSDense successfully reconstructs both the open and closed conformations from SAXS data at resolutions better than 7 Å. Furthermore, for the mixtures examined here, XSSDense provides reasonable estimates of the state populations, despite representing the conformational ensemble with a single electron density.

## Discussion

XSSDense combines experimental refinement in electron-density space with structural priors learned by a variational autoencoder (VAE), retaining sensitivity to experimental scattering data while providing finer structural detail. Unlike deterministic autoencoders [62], the VAE learns a regularised latent distribution that supports smooth interpolation between physically plausible structures [26].

The electron densities reconstructed by XSSDense provide substantially finer resolution than conventional *ab initio* SAXS shape reconstructions [24, 63]. In a benchmark of 107 proteins, DAMMIF reconstructions achieved resolutions of 13.5 Å to 52.2 Å [64]. In contrast, XSSDense achieved resolutions of 6.5 Å and 6.8 Å for the open and closed states of AdK, respectively (Fig. 5d), and 3.3 Å for crambin (Fig. 2e), exceeding the nominal resolution of the scattering data. This is possible because the structural prior exploits correlations between low- and high-resolution features arising from the highly organized internal structure of proteins.

Compared with conventional coordinate-based forward modelling [14, 65], XSS-Dense also improves the reconstruction of ensemble-averaged densities. In the crambin benchmark, forward modelling reproduced the scattering data but failed to recover the correct average density, whereas XSSDense achieved substantially higher structural fidelity (Fig. S13).

An important question is whether the reconstructed densities are dominated by the structural prior or driven by the experimental scattering data. In *As*LOV2, the same prior yields distinct densities for the compact intermediate and unfolded state (Fig. 3e), demonstrating that the experimental scattering data discriminate between different structural states. Similarly, in AdK, a single prior encompassing both open and closed conformations allows XSSDense to recover either state depending on the target scattering profile (Fig. 5b–d).

Two limitations should nevertheless be considered. First, the structural training set must adequately represent the conformational states being reconstructed (Fig. S14). Second, VAE compression introduces some loss of structural detail, particularly at higher spatial frequencies (Fig. S15 and Fig. S16), which contributes minimally to the diffuse density features within the range of the available data (0.6*/*Å), but becomes more significant at higher *q*. The recovery of distinct compact and unfolded states for *As*LOV2 and the accurate reconstruction of a mixture of two compact AdK conformations indicate that VAE-induced smoothing does not substantially compromise the ability of XSSDense to distinguish structural states at the resolution achieved here.

SAXS naturally encodes information about conformational heterogeneity, which has long been exploited to characterize i.e. polymer disorder [66, 67]. Extracting such information for proteins is more challenging because of their structural complexity, although ensemble-based approaches have demonstrated its feasibility [68, 69]. XSS-Dense extends these capabilities by refining localized conformational heterogeneity directly from structural priors and experimental scattering data. This is demonstrated for (1) continuous heterogeneity in the unfolded state of *As*LOV2 (Fig. 3e) and for (2) discrete heterogeneity in mixtures of open and closed AdK conformations, for which approximate state populations can also be recovered (Fig. 5c,e).

We note that representing a conformational ensemble by a single electron density, as in XSSDense, is an approximation. The scattering intensity of an ensemble is the population-weighted average of the individual intensities, with populations *w*_*n*_ (∑_*n*_ *w*_*n*_ = 1) and scattering amplitudes *A*_*n*_(**q**), which generally differs from the scattering calculated from the average electron density:

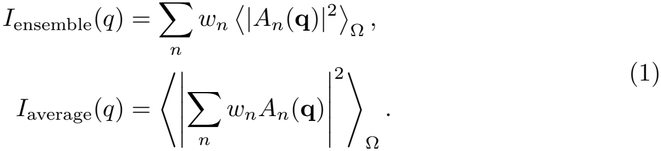

This means that scattering calculated from an averaged electron density neglects the covariance between conformations, which contributes to the ensemble scattering intensity. We quantified the resulting error for the different types of heterogeneity considered here and found it to be negligible for compact states (Fig. S17a), and small for the disordered ensemble of *As*LOV2 (Fig. S17b) and mixtures of structurally distinct AdK states (Fig. S12). An alternative would be to refine conformational ensembles, however, this risks leading to overfitting of the data. Thus, we believe that accepting the single-density approximation is justified, but care is advised in cases when the predicted structural heterogeneity becomes very large.

With respect to integrative structural biology, refinement in electron-density space has several advantages. It should become possible to include detergent micelles or nanodiscs during the refinement of membrane proteins, potentially overcoming the cumbersome treatment of detergent in all-atom refinement schemes and benefit from methods developed for single-particle cryo-electron microscopy [65, 70, 71]. XSSDense could also be integrated with current molecular simulation and structure-prediction approaches to determine conformational ensembles of intrinsically disordered proteins and RNA molecules, where structural heterogeneity is central to biological function [22, 72, 73]. Finally, additional experimental restraints in electron density space, such as NMR, electron microscopy, or electron paramagnetic resonance measurements, can be incorporated into the VAE training, extending general frameworks for combining structural priors with complementary experimental data [74].

More generally, our results establish direct electron-density refinement as a practical framework for integrating structural priors with experimental S/WAXS data. We anticipate that this representation will enable increasingly powerful integrative structural biology approaches for studying dynamic and conformationally heterogeneous biomolecular systems.

## Methods

### Generation of training and validation datasets

The structures for crambin (synthetic case) were generated from molecular dynamics simulations in a Generalized Born implicit solvent. The reference model was obtained from the RCSB Protein Data Bank (PDB ID 1CRN) [39]. The structure was minimized and later equilibrated under constant volume and temperature conditions using 2 fs time steps for 10 ns, using the molecular dynamics software NAMD and the CHARMM36 force field. The temperature was set to 310 K and the implicit solvent ion concentration to 0.3 M. The trajectory was saved every 1 ps resulting in 10 000 frames, which were subsequently aligned with the Python package MDAnalysis [75, 76]. The structures for *As*LOV2 and PYP were generated using AFsample2 with dropout enabled [37]. To ensure a large conformational landscape, the multiple sequence alignment masking was varied from 10 % to 60 % in steps of 10 % for the different data sets (Tab. S2). The resulting structures were aligned using TMalign [77]. The structures for adenylate kinase were taken from published molecular dynamics simulations of ligand-free AdK [60, 61], from which 5370 frames balanced between open-like and closed-like conformations were selected (Fig. S10). All structures were aligned on the C*α* atoms of the CORE domain (residues 1 to 29, 60 to 121 and 160 to 214) using MDAnalysis.

### Voxelisation of the electron density

The electron density of the proteins in vacuum was computed based on elastic scattering form factors from Su and Coppens (1997) [78], which fit *f* (*q*) to a sum of 6 Gaussians of the form

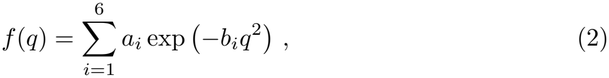

where *q* = 4*π* sin(*θ*)*/λ* is the scattering vector, and *a*_*i*_ and *b*_*i*_ are fitting parameters for each element. The real space electron density, related by the inverse Fourier transform for the atomic scattering factors [79] summed over all *N* atoms in the protein, takes the form

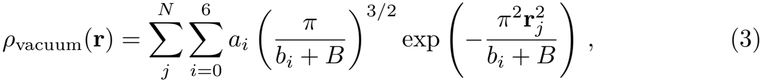

where the *B* = 8*π*^2^ (*dv/*2)^2^ factor introduces a smearing that mimics vibrational motion of the atoms. The electron density was evaluated over a 52 × 52 × 52 grid with a spacing *dv* that was system-dependent to keep the grid number constant. For the crambin the grid spacing was 1.61 Å, for *As*LOV2 2.40 Å, for PYP 2.16 Å, and for AdK 2.40 Å (Fig. S9).

### X-ray scattering calculations

The scattering amplitude of the protein corresponds to the three-dimensional Fourier transform of its voxelised electron density

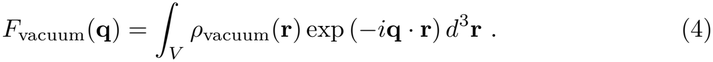

We modelled the solution scattering signal by subtracting the displaced solvent and adding a hydration-shell contrast term. To establish the protein-solvent boundary, the we defined the excluded-volume mask

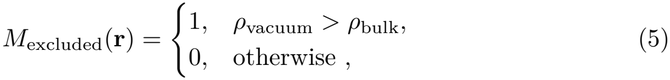

where *ρ*_bulk_ = 0.334 e*/*Å^3^ is the density of the bulk solvent (water), which ensures solvent-subtracted density is non-negative. The binary mask was then used to calculate the scattering amplitude of the excluded volume via

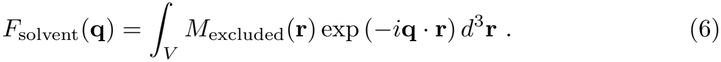

Similarly, we defined a hydration-shell mask that expanded some distance outwards from the solvent-protein boundary, thus making the hydration shell scattering amplitude contribution

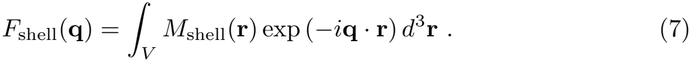

Combined, all scattering contributions yield the following expression for the total scattering amplitude

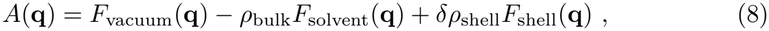

where *δρ*_shell_ is the hydration-shell contrast. Finally, we obtained one-dimensional scattering intensity profiles *I*(*q*) = ⟨|*A*(**q**)|^2^⟩_Ω_, where ⟨⟩_Ω_ represents spherical averaging. For all scattering calculations, we apply zero padding to the voxelised electron density to avoid artifacts in the Fourier transform. To ensure accurate calculations we compared the scattering intensity from the voxelised electron density against results from PEPSI-SAXS [80] for *As*LOV2 and PYP *in vacuo* and in solvent (Fig. S6).

### Validity of single-density approximation for heterogeneous ensembles

To quantify the approximation introduced by representing a conformational ensemble with a single electron density (see Discussion), we calculated the difference between the population-weighted scattering intensities and the scattering from the corresponding averaged density. This difference is given by the variance of the scattering amplitudes,

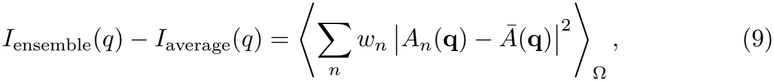

where *Ā*(**q**) = ∑*w*_*n*_*A*_*n*_(**q**).

To assess the significance of this difference for the reconstruction, we evaluated the error using the same scattering range, weighting and scaling procedures as in the XSSDense refinement, rather than comparing the scattering intensities directly. For *As*LOV2, this involves the *q*-weighted difference scattering and optimization of the excitation yield, whereas for AdK we used absolute scattering with the scale and background determined by least squares. The resulting fit-weighted errors therefore reflect the extent to which the single-density approximation limits agreement with the scattering data (Fig. S17 and Fig. S12).

### Architecture of the variational autoencoder, training and validation

The variational autoencoder (VAE) is used to generate new electron densities from a low-dimensional representation of the conformational space it has been trained on. The model, consisting of an encoder and a decoder, has the following architecture (Tab. S3 and Tab. S4).

The encoder consists of three convolutional blocks. The first two blocks each contain two 3D convolutional layers (3 × 3 × 3 kernels) followed by a max-pooling by a factor of two. The number of feature channels increases from 16 to 32 and 64. The final block contains two additional 3 × 3 × 3 convolutions and one 1 × 1 × 1 convolution. The resulting feature map is flattened and passed through a dense layer to produce a mean and log-variance 8-dimension latent distribution. The number of latent space dimensions is chosen to be approximately consistent with the number of Shannon channels in the scattering profiles (Fig. S7).

The decoder closely follows the encoder. First, the latent vector is mapped through a dense layer and reshaped. This is followed by blocks of 3D convolutional layers separated by upsampling with a factor of two. A final 3D convolutional layer with a softplus activation function ensures a positive density, and an electron conservation layer rescales the density so that its sum equals the normalized density.

The latent space is regularised toward a normal distribution *N* (0, 1) (independent in each latent dimension) through a Kullback–Leibler (KL) divergence term [81]. The network is trained by minimizing

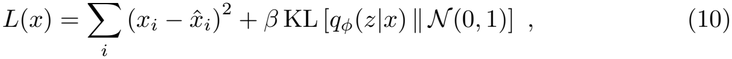

where the first term is the squared error between the input density *x* and its reconstruction 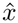, decoded from a latent vector sampled from *q*_*ϕ*_(*z*|*x*), summed over all voxels *i*, and *β* sets the strength of the regularization.

For training and validation of the variational autoencoder for all systems investigated, we first rescale the data to [0, 1] with min-max normalisation. We split the data into 90 % training and 10 % test set. The training set is further split into a 90 % training and 10 % validation set. In molecular dynamics, consecutive frames are correlated. For this reason the split is done in blocks of 100 consecutive frames, with every tenth block forming the test (or validation) dataset.

### Reconstruction algorithm

#### Difference scattering

The reconstruction algorithm is adapted from He et al. (2020) [62]. For crambin, *As*LOV2 and PYP, we applied it to the fitting of the difference scattering. The scattering from candidate densities generated with the VAE is compared against experimental data and the candidates are ranked by score based on the quality of the fit.

Two scaling steps are performed. First, we find the absolute scale between the calculated and experimental resting-state scattering by least squares,

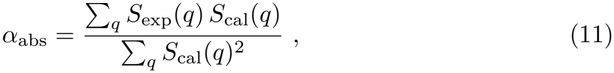

where *S*_exp_(*q*) is the experimental scattering of the resting state and *S*_cal_(*q*) is the scattering calculated from the resting-state density after encoding and decoding, such that the VAE reconstruction error cancels in the difference signal. When a solvent model is used, *α*_abs_ is fitted for each hydration shell thickness 1 Å to 5 Å and contrast 0 % to 13 % of the bulk density in steps of 1 Å and 1 %, respectively. The combination with the smallest residual ∑_*q*_ (*S*_exp_(*q*) *− α*_abs_*S*_cal_(*q*))^2^ is used for all subsequent calculations and it is reused for the fitting of the light states.

The second scaling factor corresponds to the photoactivation yield, restricted to [0, 1], and it is fitted to the *q*-weighted curves

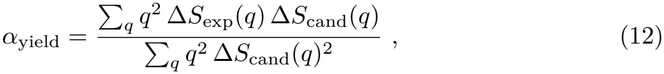

where Δ*S*_cand_(*q*) = *α*_abs_ [*S*_cand_(*q*) − *S*_cal_(*q*)] is the calculated difference scattering and *S*_cand_ is the scattering from a candidate electron density decoded from a point in the VAE latent space.

The quality of the difference fit is then evaluated with a *q*-weighted *R*^2^ [65],

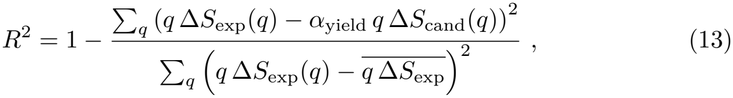

where 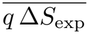 is the arithmetic mean of the *q*-weighted experimental difference scattering. Here, a value of one corresponds to a perfect fit.

Finally, to find solutions that are consistent with a prescribed photoactivation yield *α*_target_, the score includes a penalty with weight *w*,

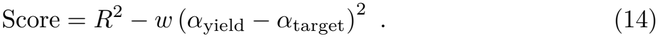

The search for candidate electron densities is performed in the latent space of the VAE with a genetic algorithm. Before the search, a pool of latent vectors is selected from the training densities such that they evenly occupy the latent space. An initial candidate population is drawn from this pool. For each generation, the best 10 % of the candidates remain unchanged, another 10 % are replaced with new candidates drawn from the pool, and the remaining candidates are crossed over and then mutated. For the crossover, each parent is paired with a random partner to produce a child 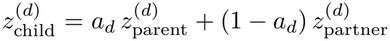, where *a*_*d*_ ∼ *U* (0, 1) is drawn separately for each dimension. In addition, each dimension from the child vector has 50 % chance of 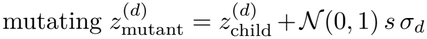, where *s* is a step size in [0.03, 1] and *σ*_*d*_ is the spread of the latent space in that dimension. The search stops when the best score has not improved for 25 generations, and the top five candidates are reported.

#### Absolute scattering

For the static AdK profiles, the absolute SAXS intensity was fitted instead of the difference scattering. Since these profiles represent stable protein conformations, the population yield is assumed to be 100 %, and the two scaling steps described above were replaced by a direct least-squares fit of the theoretical intensity to the target profile. The agreement was evaluated using

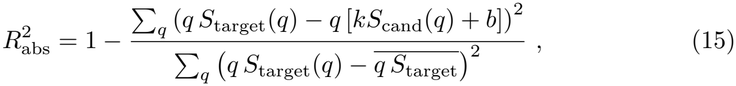

where *S*_target_(*q*) is the target SAXS intensity, *S*_cand_(*q*) is the calculated intensity of the reconstructed electron density, and 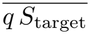 is the arithmetic mean of the *q*-weighted target intensity. The scale factor *k* and constant background *b* were determined by least squares on the *q*-weighted intensities for each candidate. The search is done with the same genetic algorithm as in the difference scattering with 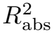 as the score and without an additional yield penalty.

### Additional methods information regarding the AdK case

Target SAXS profiles were calculated from the open (PDB ID 4AKE) and closed (PDB ID 1AKE) crystal structures of adenylate kinase (AdK), using the same scattering forward model as for the reconstructions. Profiles representing mixtures of the two conformations were generated as population-weighted sums of the open and closed scattering intensities.

To estimate the closed-state population in the reconstructed electron densities, the low-pass filtered reconstruction was decomposed into a linear combination of the lowpass filtered open and closed crystal structure densities *ρ*_cal_ ≈ *c*_open_*ρ*_open_+*c*_closed_*ρ*_closed_ using non-negative least squares. The closed-state population was then estimated as

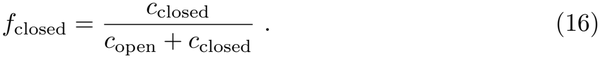

### Real space Gaussian low-pass filter

The electron densities used for training the autoencoder have a resolution determined by their grid spacing, which sets the highest representable frequency, corresponding in reciprocal space to a maximum value of *π/dv*, and in linear spatial frequency to 1*/* (2 *dv*). We apply a real-space Gaussian low-pass filter on the electron densities to suppress frequency components. For all systems, the filter has a standard deviation of 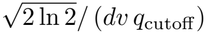 so that the signal amplitude is halved at *q*_cutoff_ = 0.6*/*Å.

## Data availability

The generated *As*LOV2 structures, the trained XSSDense model, and the unfoldedcase reconstructions are available from Zenodo: https://doi.org/10.5281/zenodo.21915224. The code required to reproduce the analyses is available at https://github.com/Westenhoff-Lab/XSSDense.

## Code availability

The source code implementing XSSDense, together with the scripts required to reproduce the experiments and analyses presented in this work, is available in the GitHub repository: https://github.com/Westenhoff-Lab/XSSDense.

## Acknowledgements

The authors acknowledge support from by H & K Jacobssons foundation. All computations were enabled by resources provided by the National Academic Infrastructure for Supercomputing in Sweden (NAISS), partially funded by the Swedish Research Council through grant agreement no. 2022-06725. The authors thank Dr. Yat Kei Lo for helpful discussions. We thank Jocelyne Vreede for providing the MD structures ensemble for PYP [52].

---

**Table S1.** System specific hyperparameters used by XSSDense, which include the VAE *β*, the target photoactivation yield and its penalty weight, and the fitted thickness and electron-density contrast of the solvation shell. Both *As*LOV2 and PYP training ensembles contain large conformational changes. A more strongly regularised latent space (larger *β*) was used to let the genetic algorithm smoothly navigate the space. For crambin, the conformational changes are subtle and required less regularisation. AdK’s open–closed hinge motion lies in the middle and using *β* = 1 or *β* = 0.01 gave qualitatively similar results that fitted the data equally well.

| System | VAE-β | Target Yield | Yield weight | Shell thickness (Å) | Shell contrast (e/Å<sup>3</sup>) |
|---|---|---|---|---|---|
| Crambin | 0.01 | 1.0 | 2.0 | – | – |
| *As*LOV2 | 1.0 | 0.15 | 5.0 | 5.0 | 0.043 |
| PYP | 1.0 | 0.42 | 5.0 | 4.0 | 0.043 |
| AdK | 0.01 | 1.0 | – | 3.0 | 0.043 |

**Table S2.** AFsample2 parameters used to generate the datasets.

| Target protein | Multiple sequence alignment masking (%) | Model preset | N-struct | Number of predictions |
|---|---|---|---|---|
| *As*LOV2 | 30–60 | Monomer | 500 | 10 000 |
| PYP | 10–50 | Monomer | 500 | 12 500 |

**Table S3.**
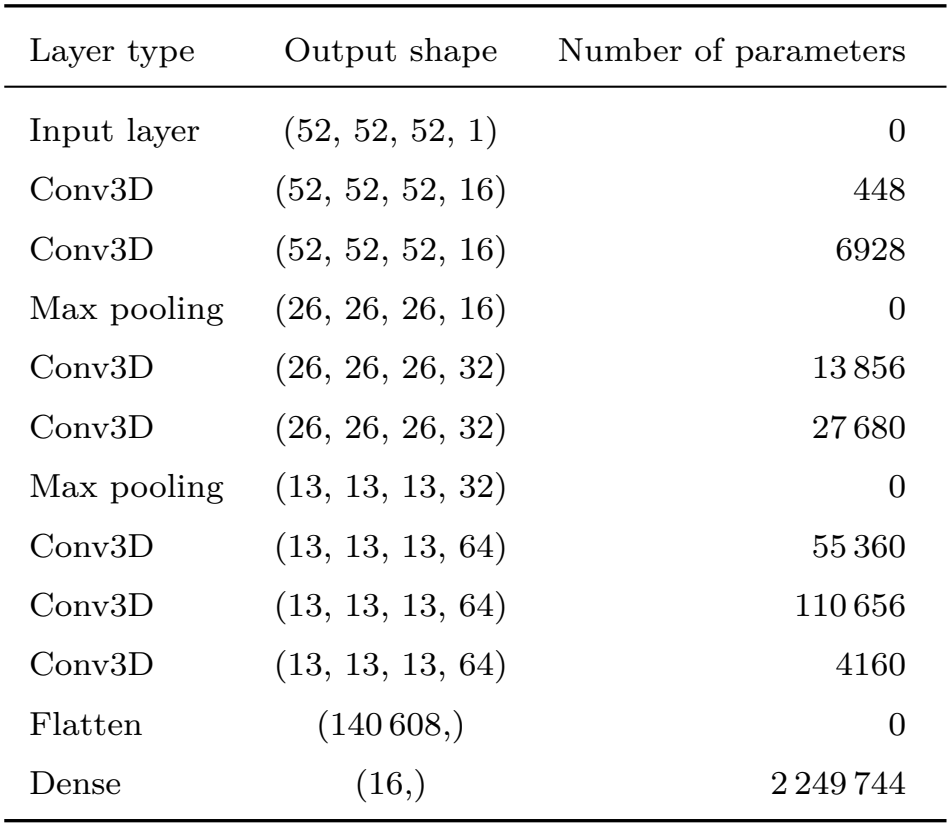
Encoder architecture. All convolutions except for the last convolution are 3 × 3 × 3 and use LeakyReLU. The last convolution is 1 × 1 × 1 and uses ReLU. Max pooling uses a factor of 2.

| Layer type | Output shape | Number of parameters |
|---|---|---|
| Input layer | (52, 52, 52, 1) | 0 |
| Conv3D | (52, 52, 52, 16) | 448 |
| Conv3D | (52, 52, 52, 16) | 6928 |
| Max pooling | (26, 26, 26, 16) | 0 |
| Conv3D | (26, 26, 26, 32) | 13 856 |
| Conv3D | (26, 26, 26, 32) | 27 680 |
| Max pooling | (13, 13, 13, 32) | 0 |
| Conv3D | (13, 13, 13, 64) | 55 360 |
| Conv3D | (13, 13, 13, 64) | 110 656 |
| Conv3D | (13, 13, 13, 64) | 4160 |
| Flatten | (140 608,) | 0 |
| Dense | (16,) | 2 249 744 |

**Table S4.** Decoder architecture. All convolutions except for the first and last convolution are 3 × 3 × 3 and use LeakyReLU. The first convolution is 1 × 1 × 1 and uses ReLU. The final convolution uses softplus so that the output electron density remains positive. Upsampling uses a factor of 2. The electron conservation rescales the grid to match the total electron count.

| Layer type | Output shape | Number of parameters |
|---|---|---|
| Input layer | (8,) | 0 |
| Dense | (140 608,) | 1 265 472 |
| Reshape | (13, 13, 13, 64) | 0 |
| Conv3D | (13, 13, 13, 64) | 4160 |
| Conv3D | (13, 13, 13, 64) | 110 656 |
| Conv3D | (13, 13, 13, 64) | 110 656 |
| Upsampling | (26, 26, 26, 64) | 0 |
| Conv3D | (26, 26, 26, 32) | 55 328 |
| Conv3D | (26, 26, 26, 32) | 27 680 |
| Upsampling | (52, 52, 52, 32) | 0 |
| Conv3D | (52, 52, 52, 16) | 13 840 |
| Conv3D | (52, 52, 52, 16) | 6928 |
| Conv3D | (52, 52, 52, 1) | 433 |
| Electron conservation | (52, 52, 52, 1) | 0 |

**Figure S1.**
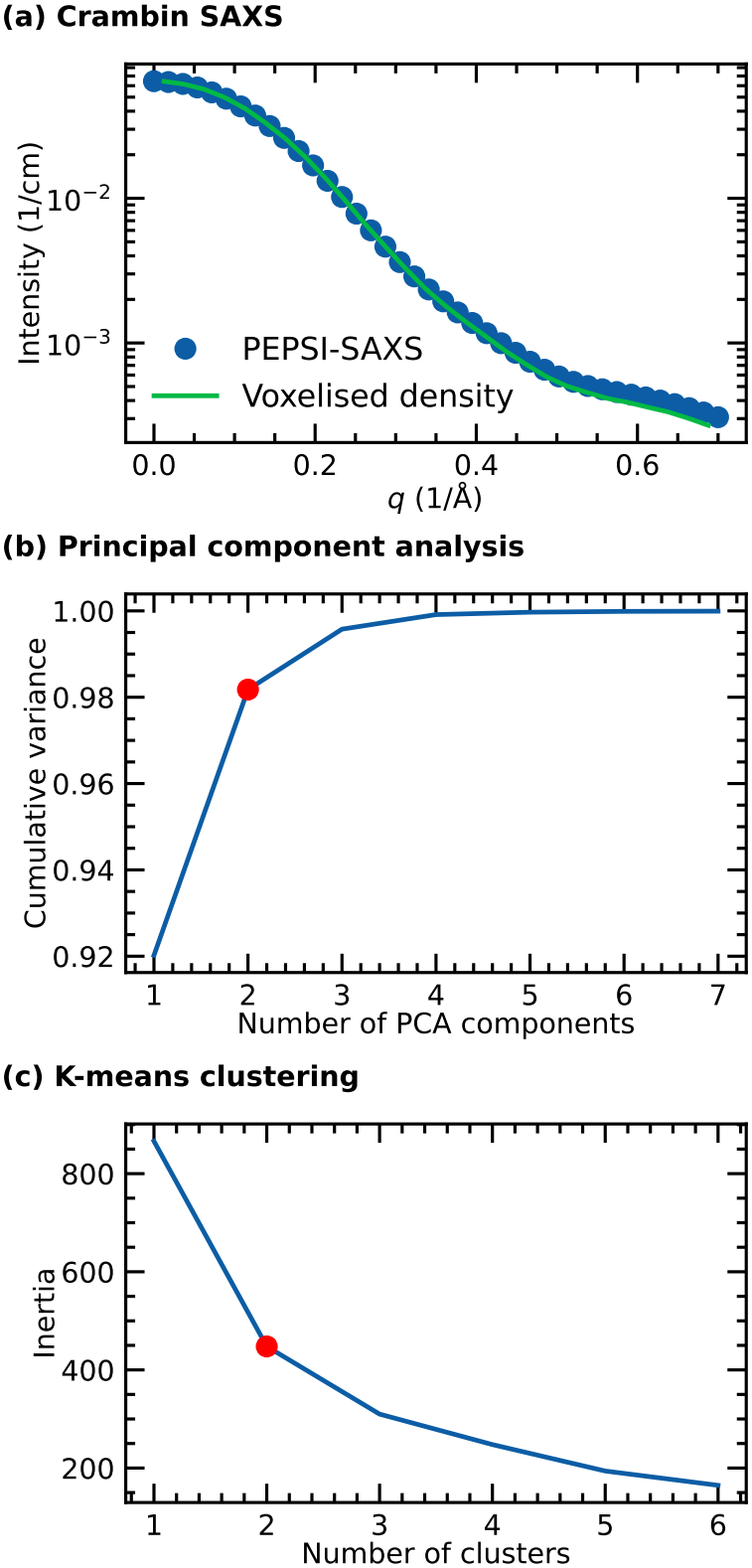
(a) Comparison of a representative small-angle X-ray scattering (SAXS) curve of crambin computed from the voxelised electron density in vacuum, against PEPSI-SAXS [80]. (b) Cumulative variance for the crambin molecular dynamics-derived SAXS dataset as a function of principal components. Using principal component analysis, two components (red circle) explain *>*95 % of the dataset variance. (c) K-means inertia (sum of the squared distances of samples to their closest cluster centre) against the number of clusters. Based on the elbow method, two clusters are optimal for classifying the low-dimensional representation of the dataset.

**Figure S2.**
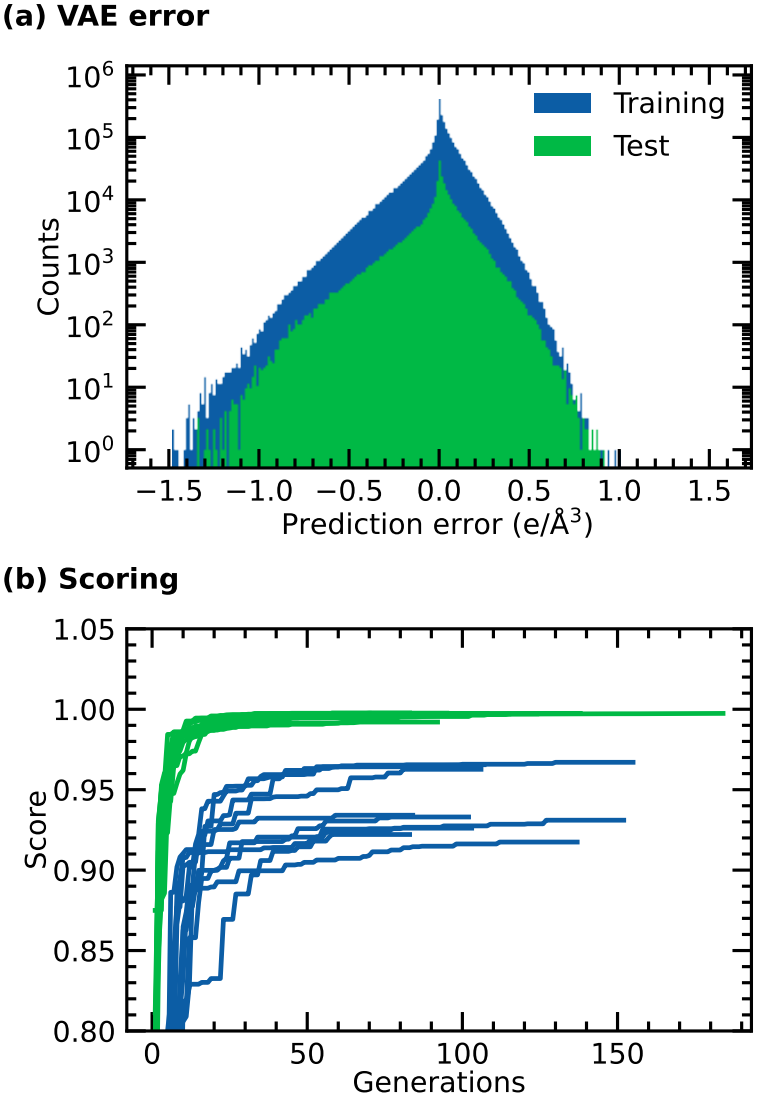
(a) Variational autoencoder prediction error of individual electron density voxels for the training and test datasets. (b) Reconstruction workflow scoring convergence, where the target *α*_yield_ is set to 1.

**Figure S3.**
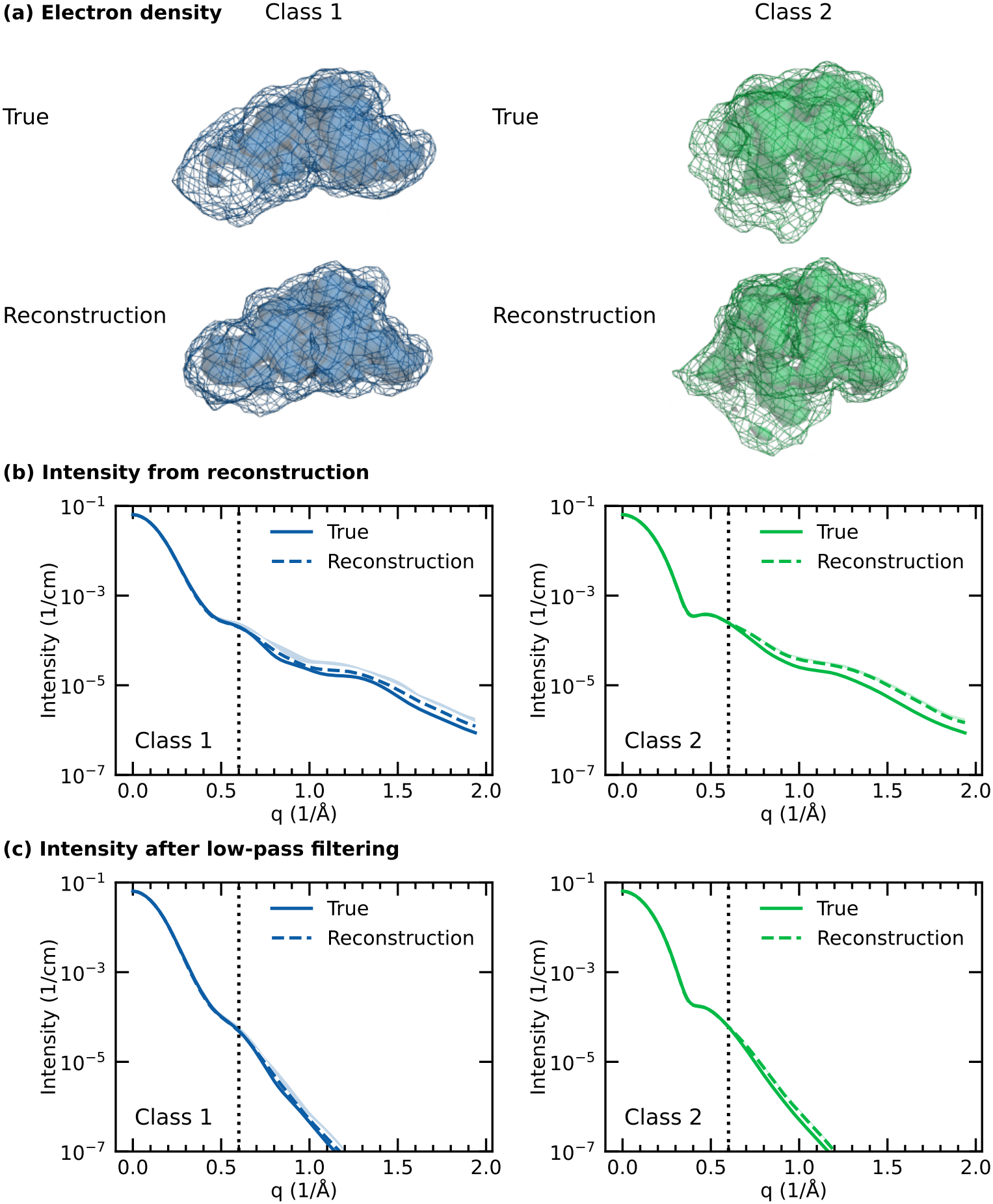
(a) Electron densities for true (MD-derived) and reconstructed (XSSDense) classes, averaged over the 10 trials. (b) Absolute scattering intensities computed from the densities in (a), for the true (solid) and reconstructed (dashed) cases. (c) Intensities computed after applying a real-space Gaussian low-pass filter with a 0.6*/*Å cut-off (dotted lines). Each panel compares (left) class 1 and (right) class 2. The mesh and surface are evaluated at 0.1 e*/*Å^3^ and 0.3 e*/*Å^3^, respectively. The light-coloured bands in (b,c) represent the min-max of the scattering over the 10 trials.

**Figure S4.**
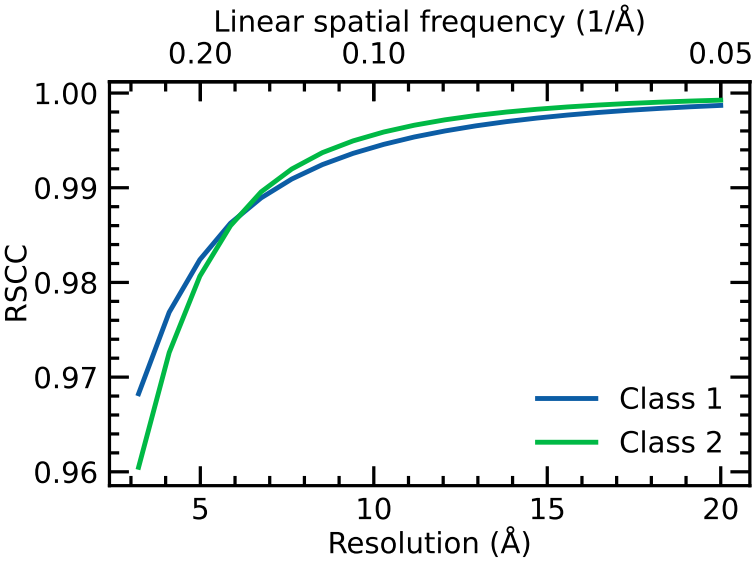
Real space Pearson correlation coefficient (RSCC) as a function of resolution comparing the true and reconstructed ensemble-averaged electron densities for two classes identified from molecular dynamics simulations of the crambin protein.

**Figure S5.**
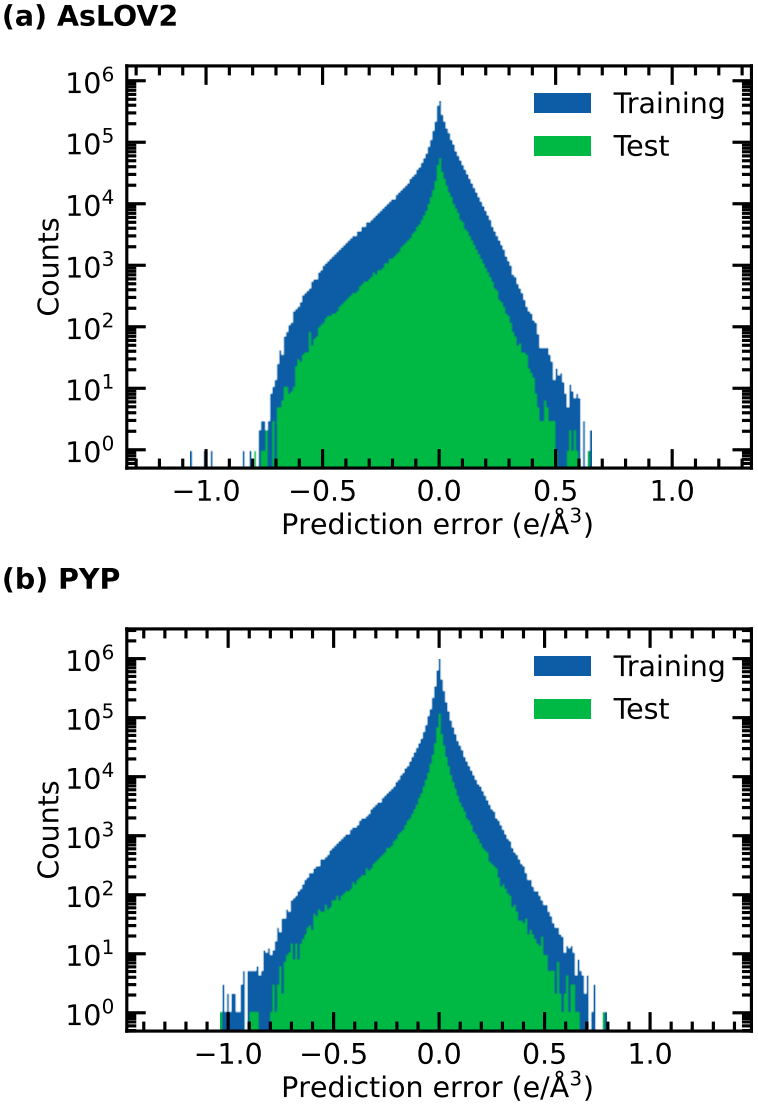
Electron density error from the variational autoencoder between the true and predicted electron densities for (a) *As*LOV2 and (b) PYP.

**Figure S6.**
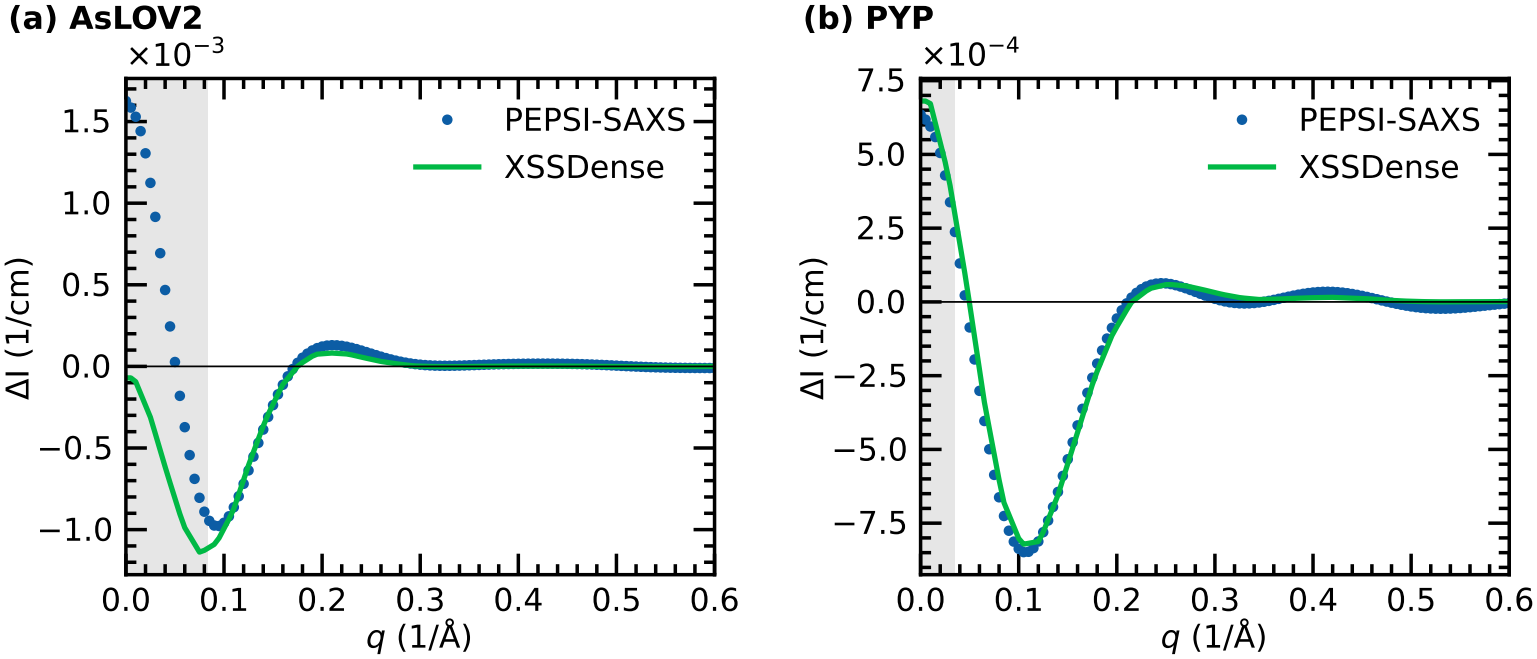
Difference scattering from voxelised electron densities compared with PEPSI-SAXS [80] for a single representative structure that is near the reconstructed (a) *As*LOV2 unfolded state and (b) the PYP pB1 state. The XSSDense curve is scaled by a fitted factor, since the magnitude of Δ*I* is absorbed by the fitted photoactivation yield. The shaded region marks the *q* below the measured data, which was not used in the reconstructions. Here, the XSSDense voxel model uses the hydration parameters from PEPSI-SAXS for both states (0.0184 e*/*Å^3^ shell contrast and 3.0 Å shell thickness of the resting structure). Differences between the curves are due to the approximate solvent mask and hydration shell of the XSSDense voxel model.

**Figure S7.**
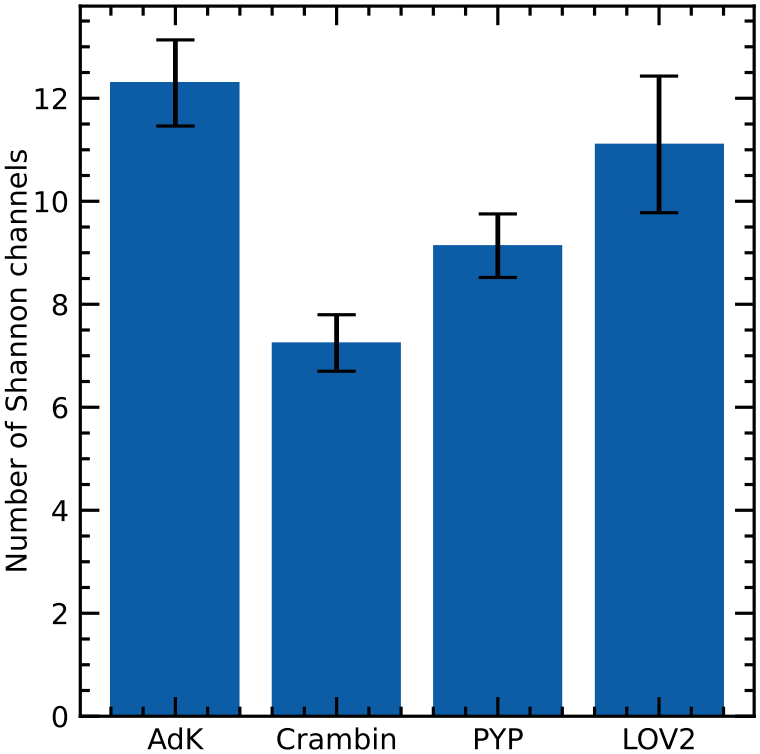
Number of Shannon channels *q*_max_*D*_max_*/π* in the scattering data. *D*_max_ was estimated from the voxelised electron density map using a threshold of 0.01 e*/*Å^3^ with *q*_max_ = 0.6*/*Å. Error bars represent the standard deviation.

**Figure S8.**
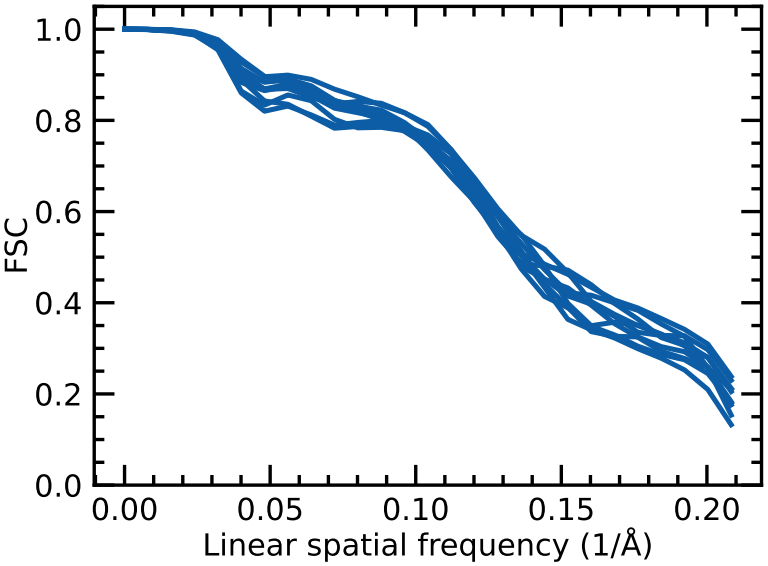
Fourier shell correlation (FSC) between the best scoring unfolded-state reconstructed density and its ten closest structures (highest FSC) in the VAE training set. The curves are nearly indistinguishable, meaning no single training density is closest to the reconstructed density.

**Figure S9.**
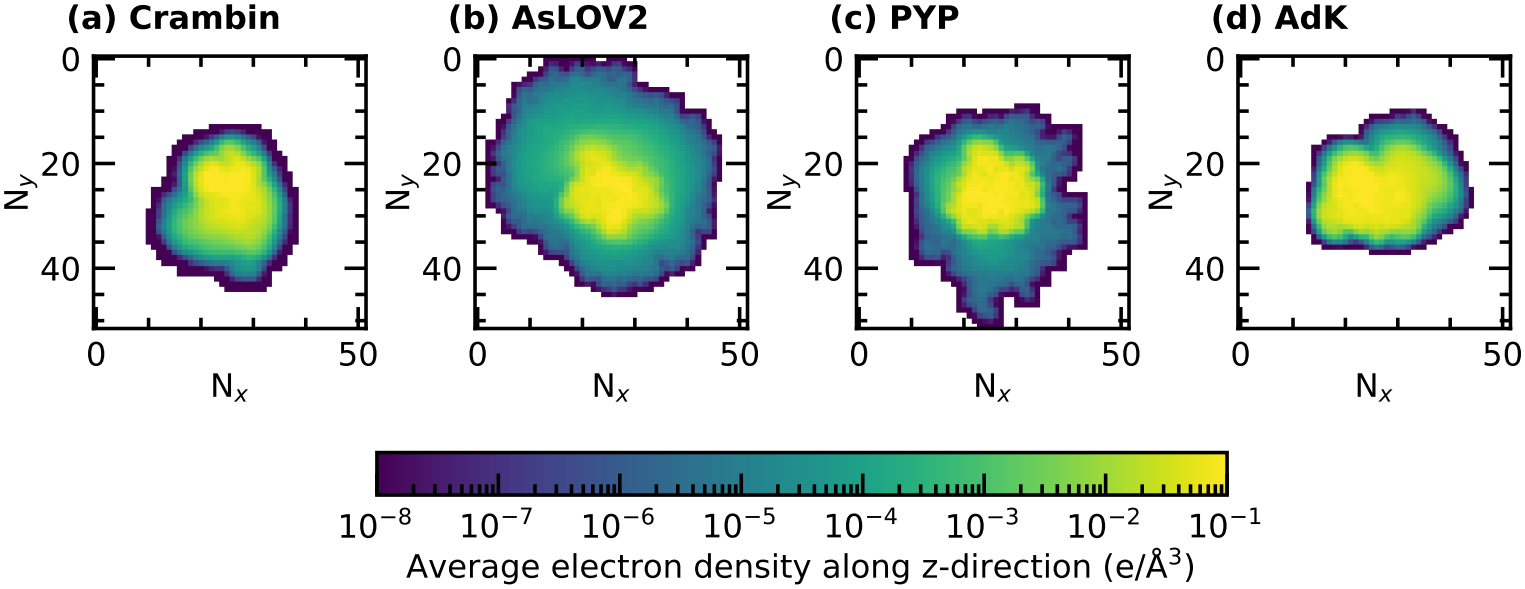
Average 52 × 52 × 52 voxelised electron density for all four proteins, projected along the z-direction. The structures are aligned prior to the voxelisation of the electron density. The areas with the highest density show regions of the protein that are most rigid, while flexible regions have a lower density. Since the rigid region appears in all training data, the autoencoder outputs a similar core region in all subsequent predicted structures.

**Figure S10.**
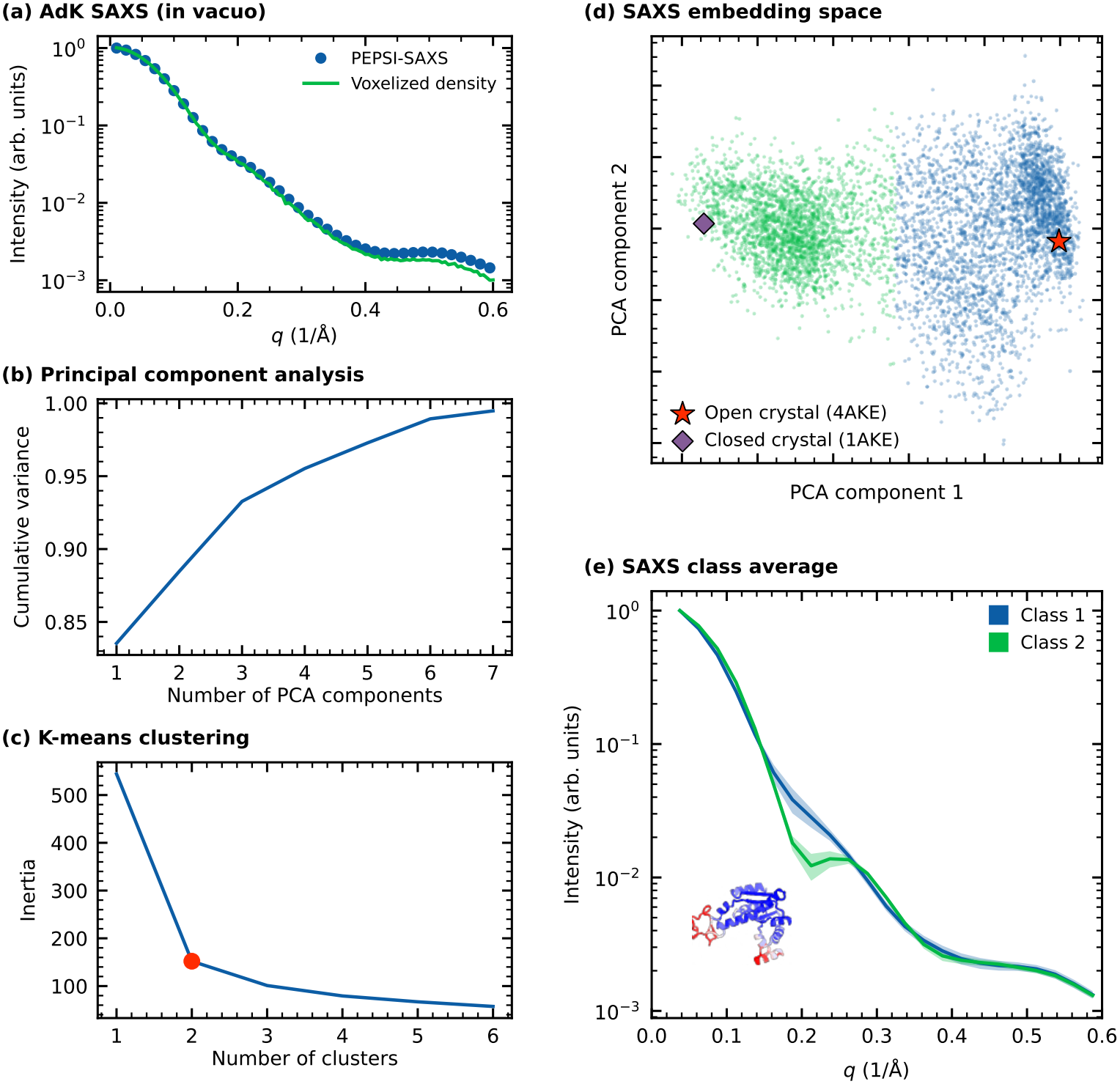
(a) Comparison of a representative small-angle X-ray scattering (SAXS) curve of adenylate kinase (AdK; open crystal structure, PDB ID 4AKE) computed from the voxelised electron density in vacuum, against PEPSI-SAXS [80]. (b) Cumulative variance for the AdK molecular dynamics-derived SAXS dataset as a function of principal components. The first component explains 83.5 % of the variance, and four components explain *>*95 %. (c) K-means inertia (sum of the squared distances of samples to their closest cluster centre) against the number of clusters. Based on the elbow method, two clusters are optimal for classifying the low-dimensional representation of the dataset. (d) Lower dimension representation and clustering of *in vacuo* SAXS profiles of AdK obtained from molecular dynamics simulations [60, 61]. The open (PDB ID 4AKE) and closed (PDB ID 1AKE) crystal structures are projected onto the same components. (e) Ensemble-average SAXS profiles (solid line) and standard deviation (shaded region) for the two identified classes. Insert shows AdK’s open crystal structure as a cartoon representation colour coded to show regions with low (blue) and high (red) root mean square fluctuation based on molecular dynamics simulations. The CORE domain is rigid, while the LID and NMP domains are flexible.

**Figure S11.**
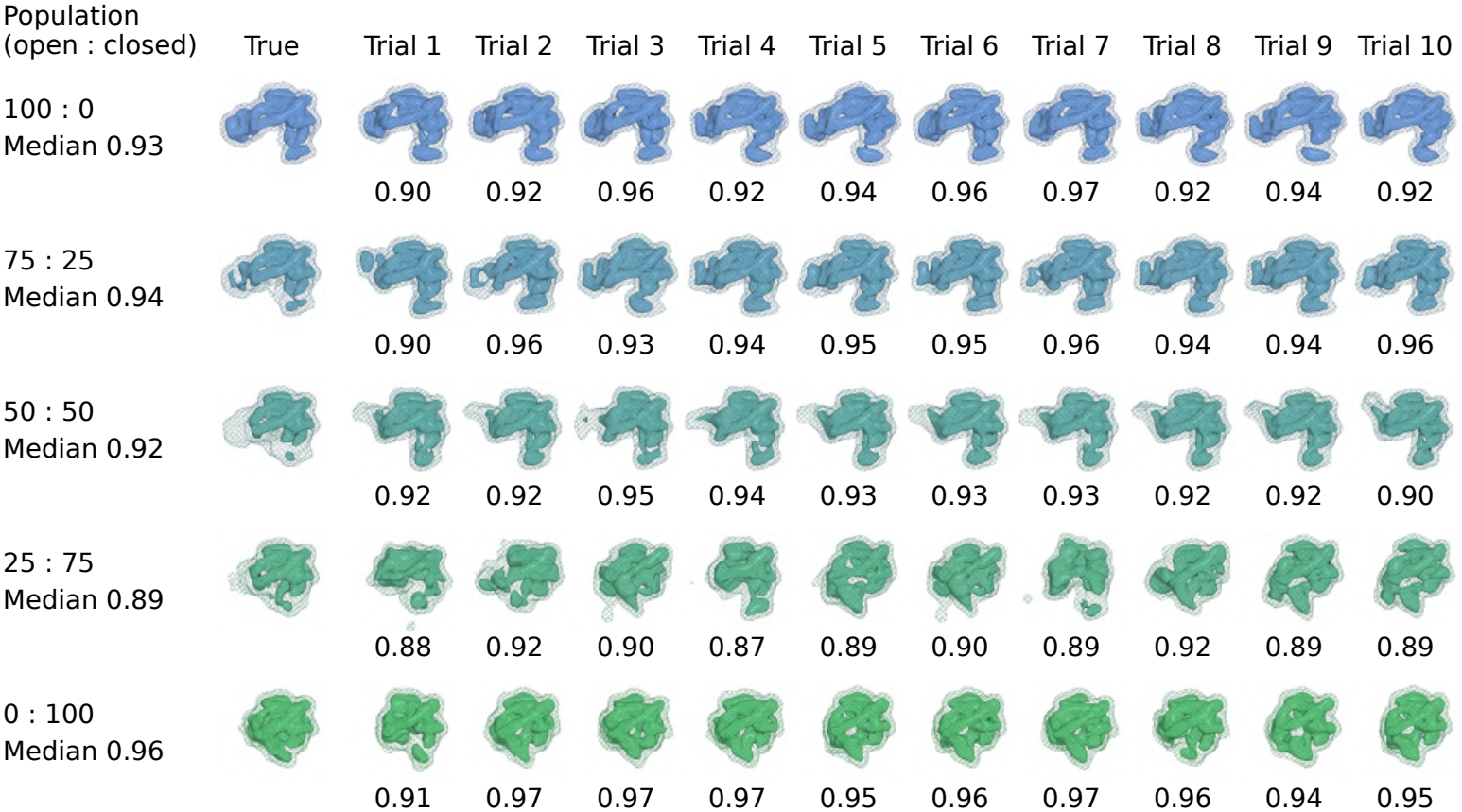
True (first column) and reconstructed electron densities of all 10 trials of adenylate kinase (AdK) at defined population ratios of the open and closed states (open : closed), plotted as a mesh at 0.1 e*/*Å^3^ and surface at 0.3 e*/*Å^3^. The maps were subjected to a low-pass Gaussian filter with a half-power point of 0.6*/*Å. Numbers below each trial give the Pearson correlation coefficient with the true electron density, and the row label gives the median of 10 trials. All trials reproduce the true density closely (median 0.89 to 0.96), so the averaged densities (Fig. 5c) represent the single reconstructions. For the pure states and the 75:25 mixture, all trials are close to the true density; for 50:50 the trials give similar shapes; and 25:75 shows the largest spread, with trials ranging from half-open to closed.

**Figure S12.**
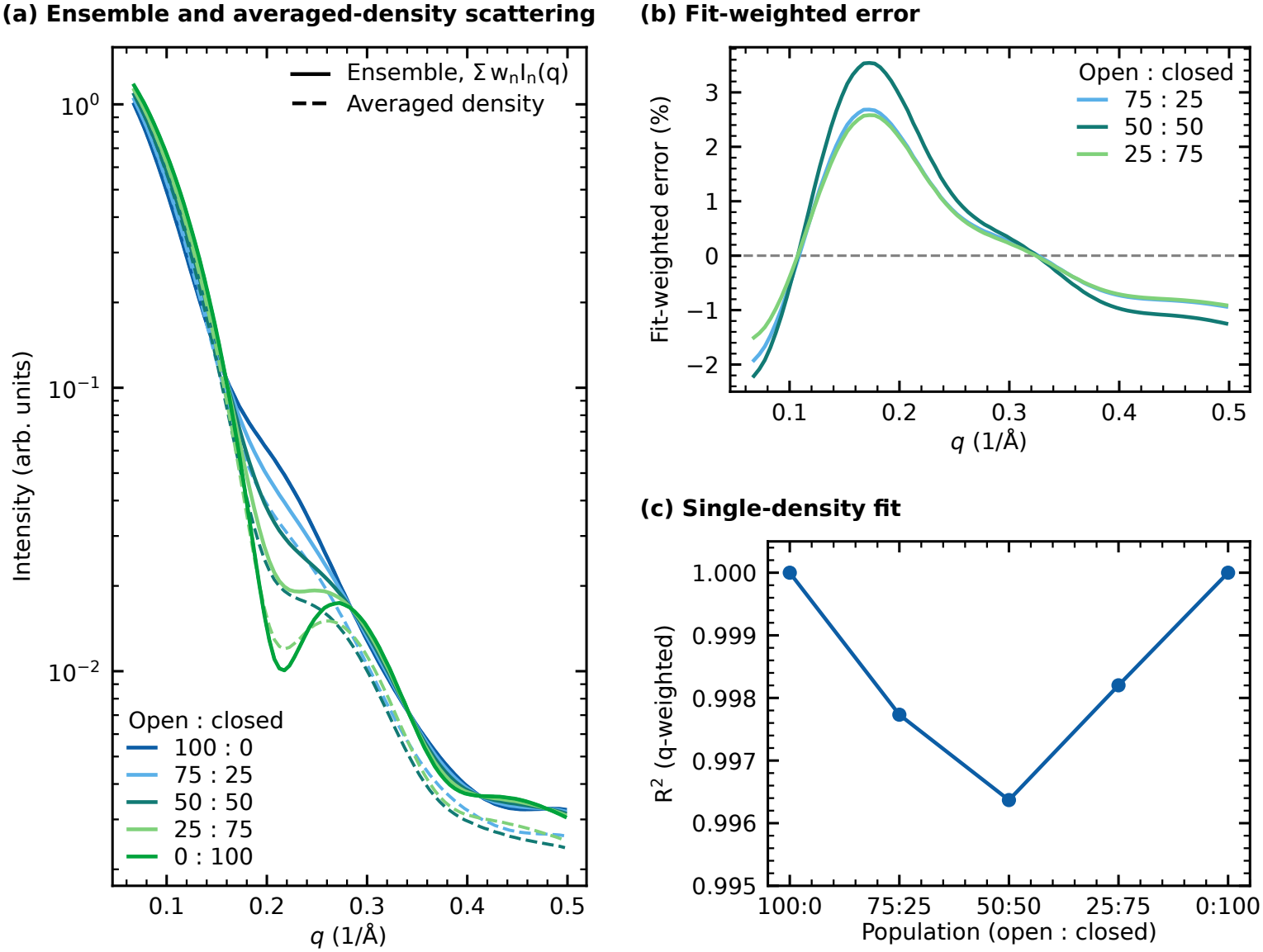
(a) Scattering of open–closed ensembles of adenylate kinase (AdK), *I*_ensemble_(*q*) = ∑_*n*_ *w*_*n*_*I*_*n*_(*q*) (solid), compared with the scattering of the population-weighted averaged electron density, *I*_average_(*q*) (dashed), for populations (open : closed) of 100:0, 75:25, 50:50, 25:75 and 0:100. The open (PDB ID 4AKE) and closed (PDB ID 1AKE) crystal densities were used. (b) Fit-weighted error, *q*[*I*_ensemble_(*q*) *−* (*kI*_average_(*q*) + *b*)], normalised to the maximum of *qI*_ensemble_(*q*). The scale *k* and constant background *b* were determined by *q*-weighted least squares, as in the reconstruction, so the error shows how much of the discrepancy between the two remains after the fit. (c) *q*-weighted *R*^2^ of the fit in (b) for each population, i.e. the best fit that a single electron density can reach. Scattering was computed *in vacuo* for *q* = 0.067*/*Å to 0.5*/*Å. The remaining error is below 4 %, and a single density reaches *R*^2^ *≥* 0.996.

**Figure S13.**
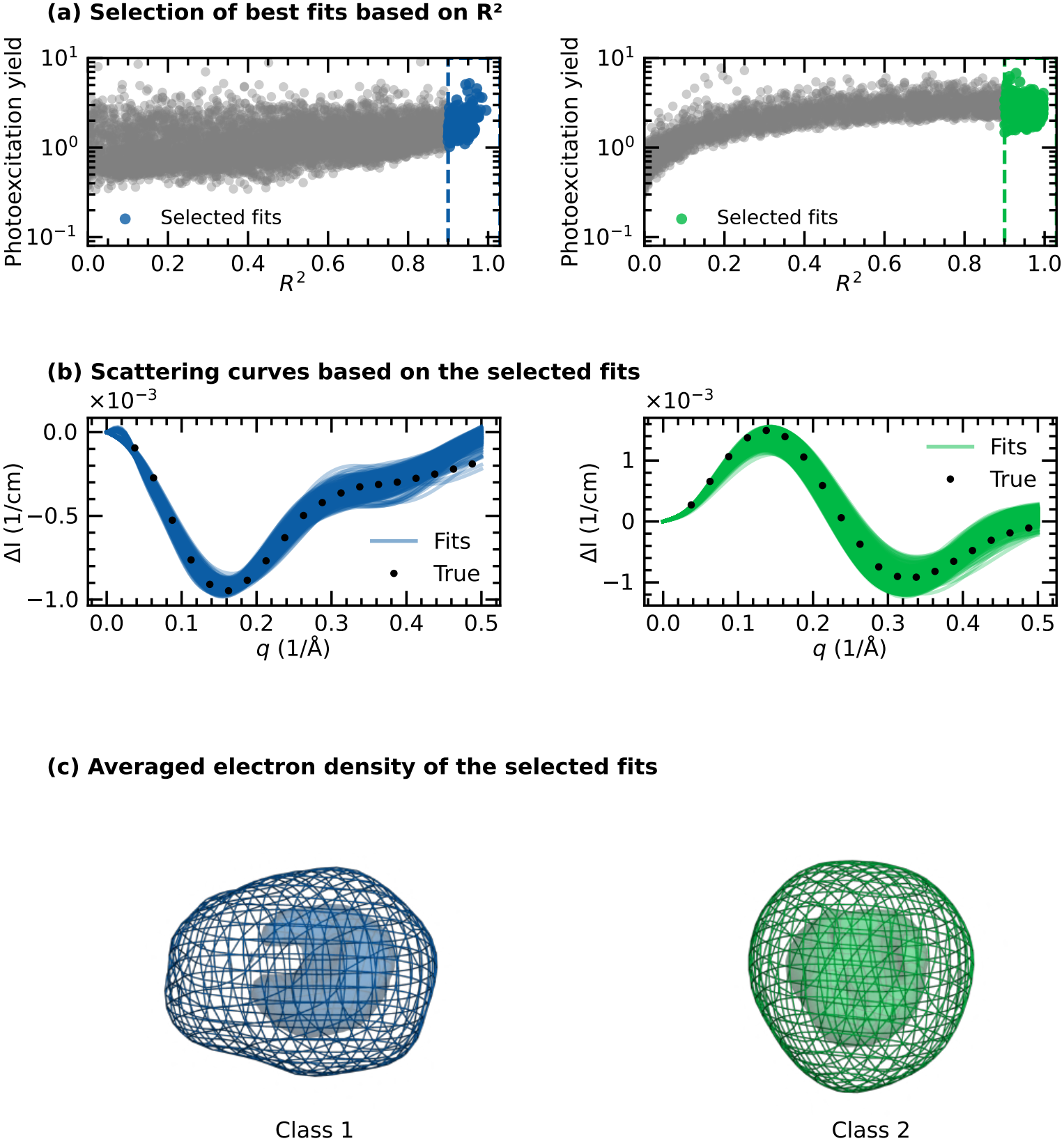
Evaluation of forward modelled scattering profiles and electron densities for crambin, performed on the difference scattering of class 1 and 2. The calculated difference profiles were fitted against the corresponding target profiles using the same MD frames as those used to train XSSDense. Scattering profiles were calculated with PEPSI-SAXS [80] using the default settings and up to *q* = 0.6*/*Å. (a) Photoexcitation yield plotted against *R*^2^ for candidate structures in class 1 (left) and class 2 (right). The photoexcitation yield was used as a scaling factor to account for discrepancies between the target and calculated scattering profiles and was not assigned a physical interpretation. Candidate structures with *R*^2^ *≥* 0.9 were selected for further analysis, yielding 234 structures for class 1 and 694 for class 2. (b) Comparison of the target and calculated difference scattering profiles for the selected candidates, with class 1 shown on the left and class 2 on the right. (c) Averaged electron densities of the selected candidates. The reconstructed densities show poor agreement with the true densities (Fig. 2d) and agreement is only evident at very low resolution.

**Figure S14.**
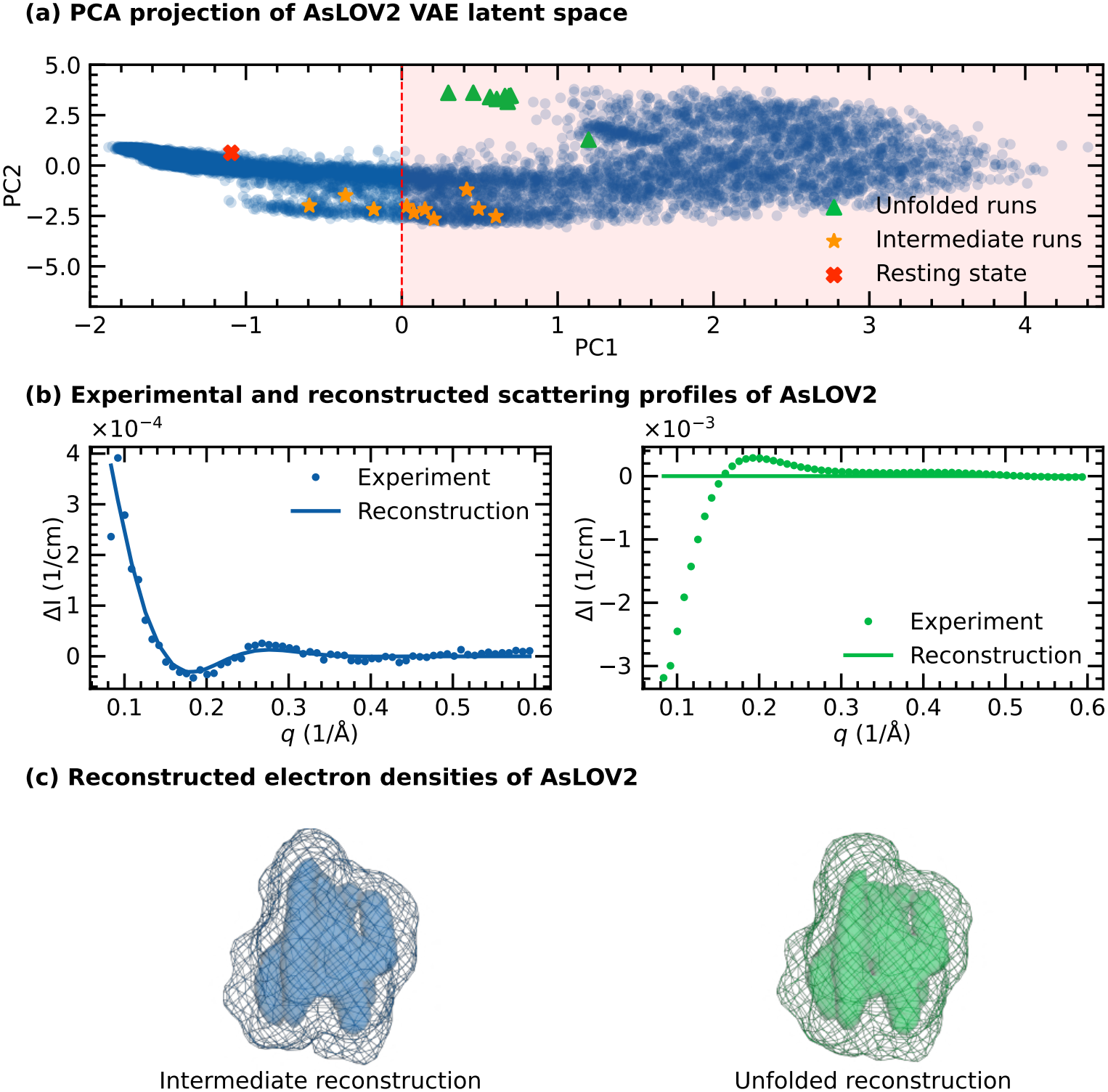
Influence of the structural prior on XSSDense reconstructions. The ground-state, intermediate-state, and unfolded-state structures were projected onto the latent space of the trained VAE to identify their locations within the learned conformational space (a). Based on this projection, a restricted subset of density maps was generated by excluding structures corresponding to the unfolded state, as well as structures in the surrounding region of the intermediate state (red shading). XSSDense was then rerun using this restricted ensemble. The intermediate state was successfully recovered, as shown by the agreement between the experimental and reconstructed scattering profiles (b, left) and the corresponding reconstructed electron density (c, left). In contrast, the unfolded state was not successfully recovered, as reflected by the disagreement between the experimental and reconstructed scattering profiles (b, right) and the corresponding electron density (c, right). These results demonstrate that the recovery of structural states depends on the conformational coverage of the structural prior and highlight the importance of including a sufficiently broad range of conformations in the training ensemble.

**Figure S15.**
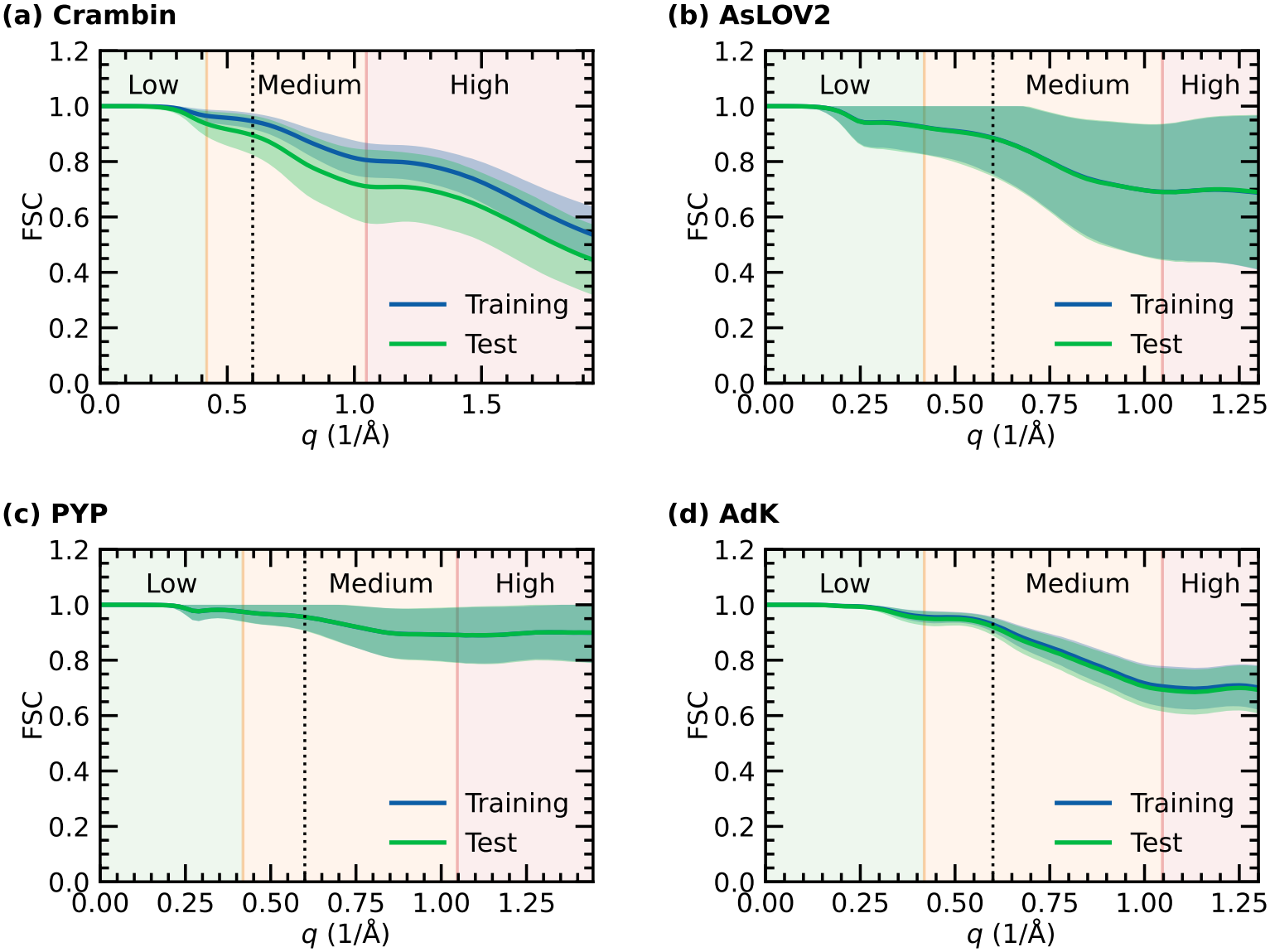
Fourier shell correlation (FSC) of all four systems investigated. Each ensemble electron density was encoded and decoded using the VAE, and the FSC was calculated between the input and reconstructed densities. The lines represent the average and shaded area the standard deviation of the train and test ensembles. The low-medium and medium-high boundaries correspond to 15 Å and 6 Å resolution, respectively. The FSC was evaluated up to the Nyquist frequency, and remains close to 1.0 showing the compression leads to minimal shift in the density within the resolution of the data (0.6*/*Å).

**Figure S16.**
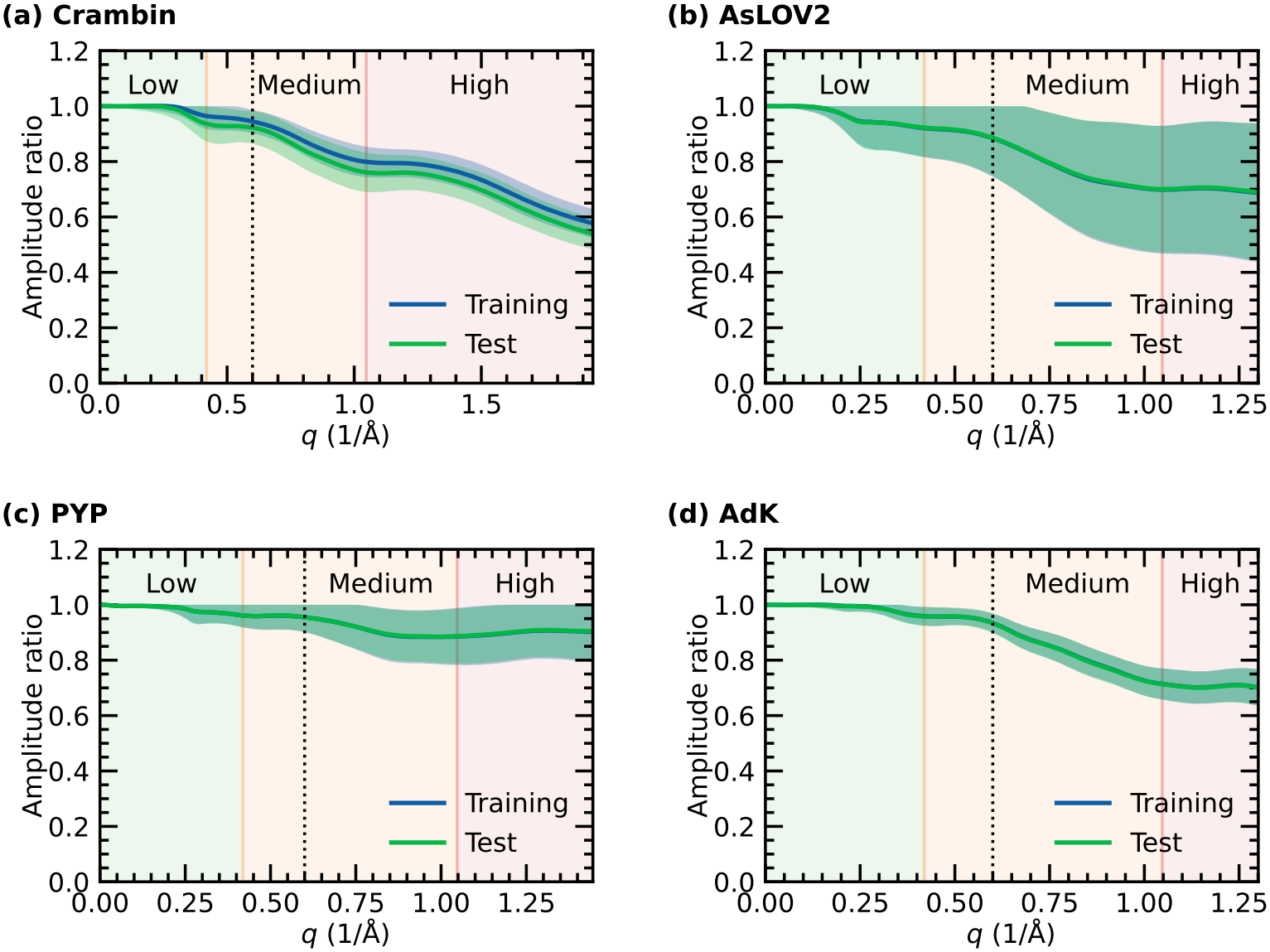
Scattering amplitude ratio 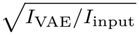 of all four systems investigated. Each ensemble electron density was encoded and decoded using the VAE, and the ratio was calculated between the input and reconstructed densities. The lines represent the average and shaded area the standard deviation of the train and test ensembles. The low-medium and medium-high boundaries correspond to 15 Å and 6 Å resolution, respectively. The ratios were evaluated up to the Nyquist frequency, and remains close to 1.0 showing the compression leads to minimal loss of structural detail within the resolution of the data (0.6*/*Å).

**Figure S17.**
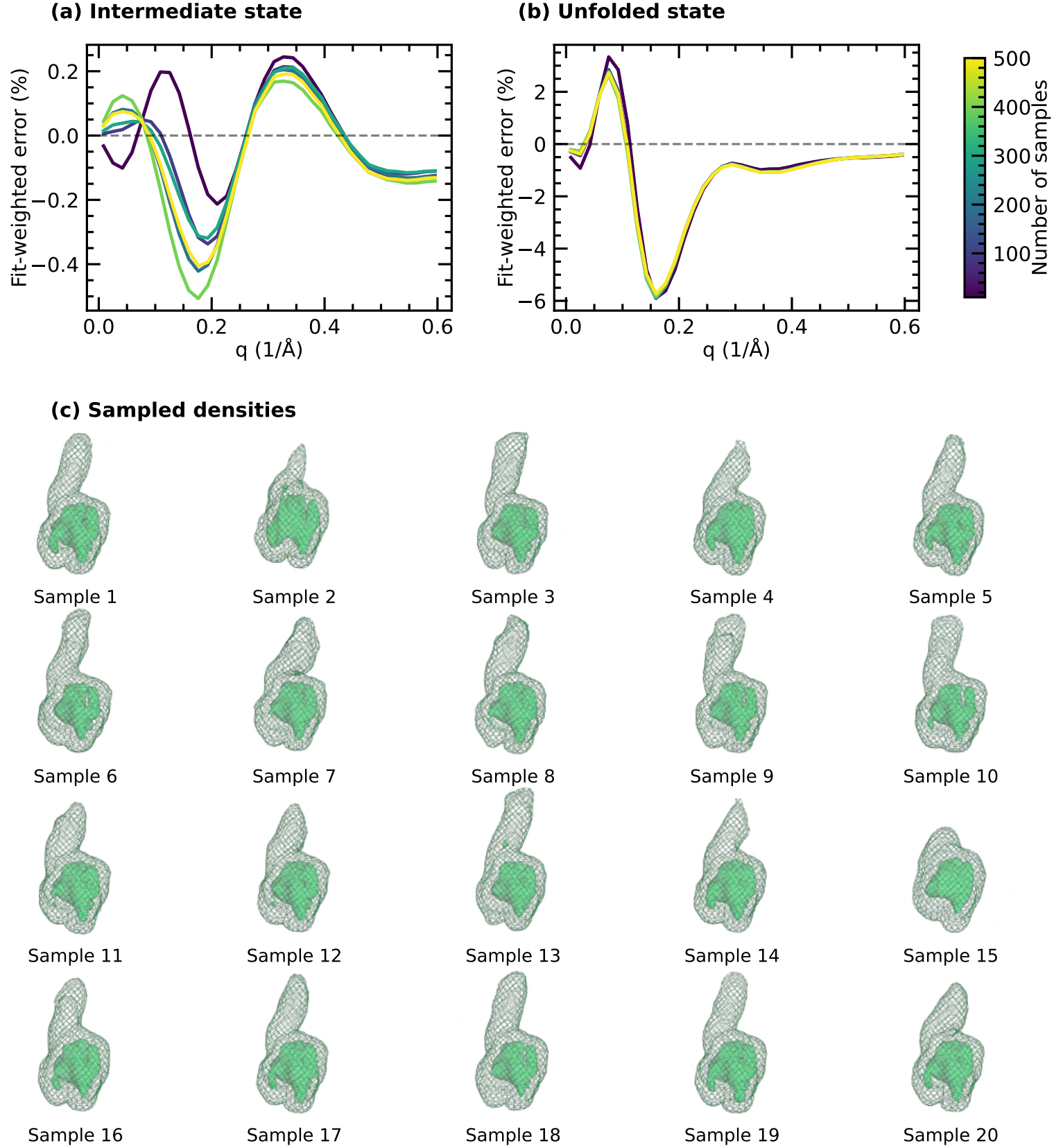
Investigating the discrepancy between the coherent and incoherent scattering for the intermediate and unfolded states of *As*LOV2. The mean (*µ*) and variance (*σ*^2^) for each dimension of the latent vector *z* were obtained by encoding the density of different states using the VAE. New latent vectors were then sampled using a Gaussian random variable *ϵ* drawn from a standard normal distribution, *N* (0, 1), resulting in sampled latent vectors *z*_sampled_ = *µ* + *σϵ*. This procedure was repeated using 10, 100, 200, 300, 400 and 500 samples. The coherent scattering calculated from the single VAE-reconstructed density, *I*_coherent_, was then compared with the incoherent scattering averaged over the sampled densities, 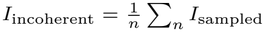. (a) Fitting error for the intermediate state, while (b) shows the fitting error for the unfolded state of *As*LOV2. In both cases, the error is *q*-weighted and normalized by the maximum value of *q I*_incoherent_. (c) Shows 20 sampled, low-passfiltered density maps of the unfolded state generated using the method described earlier. The outer meshes and inner surfaces are rendered at 0.05 e*/*Å^3^ and 0.3 e*/*Å^3^, respectively.

## Notes

https://github.com/Westenhoff-Lab/XSSDense/

### Competing Interest Statement

The authors have declared no competing interest.

### Summary of Updates

Update paper with a new case showcasing standard SAXS using AdK as a model system.

## References

[1] Henzler-Wildman, K. & Kern, D. Dynamic personalities of proteins. Nature 450, 964–972 (2007).

[2] Dmitri I Svergun & Michel H J Koch. Small-angle scattering studies of biological macromolecules in solution. Reports on Progress in Physics 66, 1735 (2003).

[3] Riback, J. A. et al. Innovative scattering analysis shows that hydrophobic disordered proteins are expanded in water. Science 358, 238–241 (2017).

[4] Ward, A. B., Sali, A. & Wilson, I. A. Integrative Structural Biology. Science 339, 913–915 (2013).

[5] Cammarata, M. et al. Tracking the structural dynamics of proteins in solution using time-resolved wide-angle X-ray scattering. Nature Methods 5, 881–886 (2008).

[6] Orädd, F. et al. Tracking the ATP-binding response in adenylate kinase in real time. Science Advances 7, eabi5514 (2021).

[7] Björling, A. et al. Structural photoactivation of a full-length bacterial phytochrome. Science Advances 2, e1600920 (2016).

[8] Cho, H. S. et al. Picosecond Photobiology: Watching a Signaling Protein Function in Real Time via Time-Resolved Small- and Wide-Angle X-ray Scattering. Journal of the American Chemical Society 138, 8815–8823 (2016).

[9] Konold, P. E. et al. Microsecond time-resolved X-ray scattering by utilizing MHz repetition rate at second-generation XFELs. Nature Methods 21, 1608–1611 (2024).

[10] Rios-Santacruz, R. et al. Integrated structural dynamics uncover a new B12 photoreceptor activation mode. Nature 650, 1045–1052 (2026).

[11] Kim, C. et al. Structural dynamics of protein-protein association involved in the light-induced transition of Avena sativa LOV2 protein. Nature Communications 15, 6991 (2024).

[12] Zielinski, K. A. et al. RNA structures and dynamics with Å resolution revealed by x-ray free-electron lasers. Science Advances 9, eadj3509 (2023).

[13] Grishaev, A., Tugarinov, V., Kay, L. E., Trewhella, J. & Bax, A. Refined solution structure of the 82-kda enzyme malate synthase G from joint NMR and synchrotron SAXS restraints. J. Biomol. NMR 40, 95–106 (2008).

[14] Hub, J. S. Interpreting solution x-ray scattering data using molecular simulations. Current Opinion in Structural Biology 49, 18–26 (2018). Theory and simulation • Macromolecular assemblies.

[15] Ramachandran, P. L. et al. The short-lived signaling state of the photoactive yellow protein photoreceptor revealed by combined structural probes. Journal of the American Chemical Society 133, 9395–9404 (2011). PMID: 21627157.

[16] Xu, X. et al. Dynamics in a pure encounter complex of two proteins studied by solution scattering and paramagnetic nmr spectroscopy. Journal of the American Chemical Society 130, 6395–6403 (2008). PMID: 18439013.

[17] Shevchuk, R. & Hub, J. S. Bayesian refinement of protein structures and ensembles against saxs data using molecular dynamics. PLOS Computational Biology 13, 1–27 (2017). URL 10.1371/journal.pcbi.1005800.

[18] Hermann, M. R. & Hub, J. S. Saxs-restrained ensemble simulations of intrinsically disordered proteins with commitment to the principle of maximum entropy. Journal of Chemical Theory and Computation 15, 5103–5115 (2019). PMID: 31402649.

[19] Chen, P.-c. & Hub, J. S. Validating Solution Ensembles from Molecular Dynamics Simulation by Wide-Angle X-ray Scattering Data. Biophysical Journal 107, 435–447 (2014).

[20] Ostojić, L. et al. Time-resolved x-ray solution scattering observations of light-induced structural changes in sensory rhodopsin ii. Structure 34, 496–507.e3 (2026).

[21] Linse, J.-B., Cho, H. S., Schotte, F., Anfinrud, P. A. & Hub, J. S. Depletion of the protein hydration shell with increasing temperature observed by small-angle x-ray scattering and molecular simulations. Journal of the American Chemical Society 147, 47117–47125 (2025). PMID: 41378747.

[22] Bernetti, M., Hall, K. B. & Bussi, G. Reweighting of molecular simulations with explicit-solvent saxs restraints elucidates ion-dependent rna ensembles. Nucleic Acids Research 49, e84–e84 (2021).

[23] Chamberlain, S. R., Moore, S. & Grant, T. D. Fitting high-resolution electron density maps from atomic models to solution scattering data. Biophys. J. 122, 4567–4581 (2023).

[24] Svergun, D. Restoring Low Resolution Structure of Biological Macromolecules from Solution Scattering Using Simulated Annealing. Biophysical Journal 76, 2879–2886 (1999).

[25] Grant, T. D. Ab initio electron density determination directly from solution scattering data. Nature Methods 15, 191–193 (2018).

[26] Kingma, D. P. & Welling, M. Auto-Encoding Variational Bayes (2013).

[27] Hoffmann, J. et al. Data-Driven Approach to Encoding and Decoding 3-D Crystal Structures (2019).

[28] Xiao, S., Song, Z., Tian, H. & Tao, P. Assessments of Variational Autoencoder in Protein Conformation Exploration. Journal of Computational Biophysics and Chemistry 22, 489–501 (2023).

[29] Luo, X. et al. Deep learning generative model for crystal structure prediction. npj Computational Materials 10, 254 (2024).

[30] Noh, J. et al. Inverse Design of Solid-State Materials via a Continuous Representation. Matter 1, 1370–1384 (2019).

[31] Huang, J., Jin, Y., Shi, Q. & Teng, D. From static structures to dynamic landscapes: Generative artificial intelligence for protein conformational dynamics. Current Opinion in Structural Biology 98, 103279 (2026).

[32] Zhong, E. D., Bepler, T., Berger, B. & Davis, J. H. CryoDRGN: Reconstruction of heterogeneous cryo-EM structures using neural networks. Nature Methods 18, 176–185 (2021).

[33] Abramson, J. et al. Accurate structure prediction of biomolecular interactions with AlphaFold 3. Nature 630, 493–500 (2024).

[34] Watson, J. L. et al. De novo design of protein structure and function with RFdiffusion. Nature 620, 1089–1100 (2023).

[35] Jing, B. et al. EigenFold: Generative Protein Structure Prediction with Diffusion Models (2023).

[36] Hsu, C., Fannjiang, C. & Listgarten, J. Generative models for protein structures and sequences. Nature Biotechnology 42, 196–199 (2024).

[37] Kalakoti, Y. & Wallner, B. AFsample2 predicts multiple conformations and ensembles with AlphaFold2. Commun. Biol. 8, 373 (2025).

[38] Katoch, S., Chauhan, S. S. & Kumar, V. A review on genetic algorithm: Past, present, and future. Multimedia Tools and Applications 80, 8091–8126 (2021).

[39] Teeter, M. M. Water structure of a hydrophobic protein at atomic resolution: Pentagon rings of water molecules in crystals of crambin. Proceedings of the National Academy of Sciences 81, 6014–6018 (1984).

[40] Zhu, J. et al. Photoadduct formation from the FMN singlet excited state in the LOV2 domain of chlamydomonas reinhardtii phototropin. The Journal of Physical Chemistry Letters 7, 4380–4384 (2016).

[41] Alexandre, M. T. A., Arents, J. C., van Grondelle, R., Hellingwerf, K. J. & Kennis, J. T. M. A base-catalyzed mechanism for dark state recovery in the avena sativa phototropin-1 lov2 domain. Biochemistry 46, 3129–3137 (2007). PMID: 17311415.

[42] Harper, S. M., Neil, L. C. & Gardner, K. H. Structural basis of a phototropin light switch. Science 301, 1541–1544 (2003).

[43] Konold, P. E. et al. Unfolding of the c-terminal jα helix in the LOV2 photoreceptor domain observed by time-resolved vibrational spectroscopy. The Journal of Physical Chemistry Letters 7, 3472–3476 (2016).

[44] Eitoku, T., Nakasone, Y., Matsuoka, D., Tokutomi, S. & Terazima, M. Conformational dynamics of phototropin 2 lov2 domain with the linker upon photoexcitation. Journal of the American Chemical Society 127, 13238–13244 (2005). PMID: 16173753.

[45] Freddolino, P. L., Gardner, K. H. & Schulten, K. Signaling mechanisms of lov domains: new insights from molecular dynamics studies. Photochem. Photobiol. Sci. 12, 1158–1170 (2013).

[46] Iuliano, J. N. et al. Unraveling the mechanism of a lov domain optogenetic sensor: A glutamine lever induces unfolding of the jα helix. ACS Chemical Biology 15, 2752–2765 (2020). PMID: 32880430.

[47] Strickland, D., Moffat, K. & Sosnick, T. R. Light-activated dna binding in a designed allosteric protein. Proceedings of the National Academy of Sciences 105, 10709–10714 (2008).

[48] Sprenger, W. W., Hoff, W. D., Armitage, J. P. & Hellingwerf, K. J. The eubacterium Ectothiorhodospira halophila is negatively phototactic, with a wavelength dependence that fits the absorption spectrum of the photoactive yellow protein. Journal of bacteriology 175, 3096–3104 (1993).

[49] Tenboer, J. et al. Time-resolved serial crystallography captures high-resolution intermediates of photoactive yellow protein. Science 346, 1242–1246 (2014).

[50] Pande, K. et al. Femtosecond structural dynamics drives the trans/cis isomerization in photoactive yellow protein. Science 352, 725–729 (2016).

[51] Konold, P. E. et al. Confinement in crystal lattice alters entire photocycle pathway of the photoactive yellow protein. Nat. Commun. 11, 4248 (2020).

[52] A. Rohrdanz M., Zheng, W., Lambeth, B., Vreede, J. & Clementi, C. Multiscale approach to the determination of the photoactive yellow protein signaling state ensemble. PLOS Computational Biology 10, 1–10 (2014).

[53] Hoff, W. D. et al. Thiol ester-linked p-coumaric acid as a new photoactive pros-thetic group in a protein with rhodopsin-like photochemistry. Biochemistry 33, 13959–13962 (1994). PMID: 7947803.

[54] Imamoto, Y., Kamikubo, H., Harigai, M., Shimizu, N. & Kataoka, M. Light-induced global conformational change of photoactive yellow protein in solution. Biochemistry 41, 13595–13601 (2002).

[55] Vreede, J., Juraszek, J. & Bolhuis, P. G. Predicting the reaction coordinates of millisecond light-induced conformational changes in photoactive yellow protein. Proceedings of the National Academy of Sciences 107, 2397–2402 (2010).

[56] Pandey, S. et al. Time-resolved serial femtosecond crystallography at the European XFEL. Nature Methods 17, 73–78 (2020).

[57] Müller, C. W., Schlauderer, G. J., Reinstein, J. & Schulz, G. E. Adenylate kinase motions during catalysis: An energetic counterweight balancing substrate binding. Structure 4, 147–156 (1996). PMID: 8805521.

[58] Henzler-Wildman, K. A. et al. Intrinsic motions along an enzymatic reaction trajectory. Nature 450, 838–844 (2007).

[59] Müller, C. W. & Schulz, G. E. Structure of the complex between adenylate kinase from Escherichia coli and the inhibitor Ap5A refined at 1.9 Å resolution. A model for a catalytic transition state. Journal of Molecular Biology 224, 159–177 (1992). PMID: 1548697.

[60] Seyler, S. & Beckstein, O. Molecular dynamics trajectory for benchmarking MDAnalysis (2017).

[61] Magkakis, K. et al. Addressing structural heterogeneity in real-time tracking of protein dynamics triggered by caged compounds. Journal of Molecular Biology 437, 169476 (2025). PMID: 41061951. Simulation data: doi:10.5281/zenodo.15853696.

[62] He, H., Liu, C. & Liu, H. Model Reconstruction from Small-Angle X-Ray Scattering Data Using Deep Learning Methods. iScience 23, 100906 (2020).

[63] Franke, D. & Svergun, D. I. DAMMIF, a program for rapid ab-initio shape determination in small-angle scattering. Journal of Applied Crystallography 42, 342–346 (2009).

[64] Tuukkanen, A. T., Kleywegt, G. J. & Svergun, D. I. Resolution of ab initio shapes determined from small-angle scattering. IUCrJ 3, 440–447 (2016).

[65] Andersson, M. et al. Structural Dynamics of Light-Driven Proton Pumps. Structure 17, 1265–1275 (2009).

[66] Debye, P. Molecular-weight determination by light scattering. The Journal of Physical and Colloid Chemistry 51, 18–32 (1947).

[67] Pedersen, J. S. Analysis of small-angle scattering data from colloids and polymer solutions: modeling and least-squares fitting. Advances in Colloid and Interface Science 70, 171–210 (1997).

[68] Bernadó, P., Mylonas, E., Petoukhov, M. V., Blackledge, M. & Svergun, D. I. Structural characterization of flexible proteins using small-angle x-ray scattering. Journal of the American Chemical Society 129, 5656–5664 (2007).

[69] Tria, G., Mertens, H. D. T., Kachala, M. & Svergun, D. I. Advanced ensemble modelling of flexible macromolecules using x-ray solution scattering. IUCrJ 2, 207–217 (2015).

[70] Bharadwaj, A., Veerbeek, L. & Jakobi, A. J. Interactive segmentation of membrane and membrane mimic densities in cryo-EM maps. bioRxiv 2026.02.25.707988 (2026).

[71] Ivanović, M. T., Hermann, M. R., Wójcik, M., Pérez, J. & Hub, J. S. Small-angle x-ray scattering curves of detergent micelles: Effects of asymmetry, shape fluctuations, disorder, and atomic details. The Journal of Physical Chemistry Letters 11, 945–951 (2020). PMID: 31951134.

[72] Tesei, G. et al. Conformational ensembles of the human intrinsically disordered proteome. Nature 626, 897–904 (2024).

[73] Ghafouri, H. et al. Toward a unified framework for determining conformational ensembles of disordered proteins. Nat. Methods 23, 705–719 (2026).

[74] Wankowicz, S. A. & Bonomi, M. From possibility to precision in macromolecular ensemble prediction. Nature Methods 23, 1100–1108 (2026).

[75] Gowers, R. J. et al. MDAnalysis: A python package for the rapid analysis of molecular dynamics simulations. SciPy 2016 (2016).

[76] Michaud-Agrawal, N., Denning, E. J., Woolf, T. B. & Beckstein, O. MDAnalysis: A toolkit for the analysis of molecular dynamics simulations. Journal of Computational Chemistry 32, 2319–2327 (2011).

[77] Zhang, Y. & Skolnick, J. TM-align: A protein structure alignment algorithm based on the TM-score. Nucleic Acids Research 33, 2302–2309 (2005).

[78] Su, Z. & Coppens, P. Relativistic x-ray elastic scattering factors for neutral atoms z = 1-54 from multiconfiguration dirac-fock wavefunctions in the 0–12 Å−1 sin(θ/λ) range, and six-gaussian analytical expressions in the 0–6 Å−1 range. Acta Crystallographica Section A 53, 749–762 (1997).

[79] Navaza, J. On the computation of structure factors by FFT techniques. Acta Crystallographica Section A 58, 568–573 (2002).

[80] Grudinin, S., Garkavenko, M. & Kazennov, A. Pepsi-SAXS : An adaptive method for rapid and accurate computation of small-angle X-ray scattering profiles. Acta Crystallographica Section D 73, 449–464 (2017).

[81] Kingma, D. P. & Welling, M. An Introduction to Variational Autoencoders. Foundations and Trends in Machine Learning 12, 307–392 (2019).

